# Interphase chromosome conformation is specified by distinct folding programs inherited via mitotic chromosomes or through the cytoplasm

**DOI:** 10.1101/2024.09.16.613305

**Authors:** Allana Schooley, Sergey V. Venev, Vasilisa Aksenova, Emily Navarrete, Mary Dasso, Job Dekker

## Abstract

Identity-specific interphase chromosome conformation must be re-established each time a cell divides. To understand how interphase folding is inherited, we developed an experimental approach that physically segregates mediators of G1 folding that are intrinsic to mitotic chromosomes from cytoplasmic factors. Proteins essential for nuclear transport, RanGAP1 and Nup93, were degraded in pro-metaphase arrested DLD-1 cells to prevent the establishment of nucleo-cytoplasmic transport during mitotic exit and isolate the decondensing mitotic chromatin of G1 daughter cells from the cytoplasm. Using this approach, we discover a transient folding intermediate entirely driven by chromosome-intrinsic factors. In addition to conventional compartmental segregation, this chromosome-intrinsic folding program leads to prominent genome-scale microcompartmentalization of mitotically bookmarked and cell type-specific cis-regulatory elements. This microcompartment conformation is formed during telophase and subsequently modulated by a second folding program driven by factors inherited through the cytoplasm in G1. This nuclear import-dependent folding program includes cohesin and factors involved in transcription and RNA processing. The combined and inter-dependent action of chromosome-intrinsic and cytoplasmic inherited folding programs determines the interphase chromatin conformation as cells exit mitosis.

## Introduction

Interphase chromosome folding is specific to the transcriptional program of a given cell type (Bickmore and van Steensel 2013; Dekker and Mirny 2016; Misteli 2020; Jerkovic and Cavalli 2021). Mechanistically, interphase folding is achieved by two major processes: Loop extrusion mediated by cohesin and spatial compartmentalisation, likely driven by contact affinities (Mirny et al. 2019). The genome is segregated in the space of the nucleus into several types of “sub-compartments” that form both within (in cis) and between (in trans) chromosomes (Lieberman-Aiden et al. 2009). Heterochromatic loci cluster together, often at the nuclear periphery, and active chromatin domains associate most frequently with each other in the nuclear interior. The factors that determine compartmentalization remain poorly understood (Hildebrand and Dekker 2020; Harris et al. 2023). The distribution of histone modifications along the genome is strongly correlated with compartment formation (Lieberman-Aiden et al. 2009; Rao et al. 2014; Wang et al. 2021), and therefore it is possible that these modifications themselves display homotypic affinities (i.e., H3K9Me3 (Gibson et al. 2019). Additionally, factors and complexes that bind to histones carrying specific modifications (e.g., BRD proteins for H3K27Ac (Dey et al. 2003; Gibson et al. 2019), or HP1 proteins for H3K9Me3 (Bannister et al. 2001; Keenen et al. 2021) can mediate interactions between marked loci. As different loci are active and inactive in different cell types, with the associated changes in chromatin composition and histone modification, their spatial positioning and clustering into sub-compartments also differs between cell types.

Interphase chromosome folding is actively modulated by cohesin-dependent loop formation. The pattern of interactions generated by loop extrusion along the genome is also highly cell type-specific due to the fact that cohesin’s activity is controlled in part by the position of cis-regulatory elements. Cohesin can be loaded throughout the genome (Spracklin et al. 2023) but appears to be more efficiently recruited at open chromatin sites such as enhancers (Valton et al. 2022; Galitsyna et al. 2023; Rinzema et al. 2022). Furthermore, loop extrusion is blocked in a directional manner at CTCF-bound sites (Rao et al. 2014; de Wit et al. 2015; Guo et al. 2015; Vietri Rudan et al. 2015) and cohesin complexes are unloaded near 3’ ends of active genes (Valton et al. 2022; Busslinger et al. 2017). Together, these elements specify a genome-wide “cohesin traffic pattern” that determines where cohesin loads, where its extrusion is blocked, and where it is dissociated. These dynamics in turn give rise to the complex and identity-specific chromosome folding pattern observed by Hi-C that includes loops (e.g., CTCF-CTCF and promoter-enhancer loops), stripes, flares, insulated boundaries, and topologically associating domains (TADs) (Fudenberg et al. 2017).

During mitosis, most if not all features of cell type-specific chromosome folding, such as compartments and cohesin-mediated looping, become undetectable by Hi-C. Instead, a largely cell type-invariant compressed array of partially self-entangled loops is formed by condensin complexes, to give rise to rod-shaped mitotic chromatids (Naumova et al. 2013; Gibcus et al. 2018; Samejima et al. 2024; Dekker 2014; Hildebrand et al. 2024). These loops are not positioned at any specific cis-regulatory elements. Many of the chromatin-associated factors that contribute to interphase organization are absent from mitotic chromosomes. Condensins actively remove loop extruding cohesin complexes during prophase of mitosis (Samejima et al. 2024) and cell cycle kinases drive the phosphorylation and dissociation of a large proportion of the chromatin-bound factors, including CTCF, transcription factors and, to a large extent, RNA polymerases (Oomen et al. 2019; Sekiya et al. 2017; Martínez-Balbás et al. 1995; Parsons and Spencer 1997; Samejima et al. 2022).

However, not all regulatory factors dissociate from mitotic chromosomes, and a number remain bound at specific cis-elements. This phenomenon, referred to as mitotic bookmarking (Ito and Zaret 2022), records active loci for re-activation in the next cell cycle. Cell cycle stage-dependent changes in chromosome structure are also observed at the scale of nucleosomes. While promoters that are active in a given cell type remain largely nucleosome-free, enhancers are less accessible during mitosis and are likely bookmarked to be reopened by remodelers during the next cell cycle (Hsiung et al. 2015; Oomen et al. 2019). Similarly, CTCF sites are thought to be bookmarked by active histone modifications and become occupied by nucleosomes in mitosis, although the extent to which this occurs differs between cell types (Oomen et al. 2019, 2023; Chervova et al. 2023).

The dramatic changes to chromosome organization that occur during mitosis pose a particular challenge to cycling cells. Every time cells re-enter G1, the cell type-invariant mitotic chromosome structure must be converted to an identity-specific interphase conformation. The relative timing of events during this process has been delineated in several recent studies (Zhang et al. 2019; Abramo et al. 2019; Pelham-Webb et al. 2021; Kang et al. 2020; Brunner et al. 2024). First, condensins dissociate during telophase and a transient chromosome folding state devoid of extruded loops is formed (Abramo et al. 2019). Although CTCF is rapidly enriched at post-mitotic chromatin, the cohesin complex is recruited later during cytokinesis (Zhang et al. 2019; Abramo et al. 2019) at which point extrusion rapidly re-establishes loops, TADs, and other extrusion-related features. Genome compartmentalization is first detected around telophase, but does not reach the full extent of segregation until well into G1 (Abramo et al. 2019) .

Although the kinetics of post-mitotic chromosome re-organization are increasingly known, it remains unclear how the cell type-specific folding program is transmitted through mitosis. Observations of mitotic bookmarking by histone modifications and retained factors at cis elements, as well as maintained accessibility at active promoters, suggests that at least part of the folding program is carried on the mitotic chromosomes. These factors must be inactive or overruled by condensins during mitosis and then re-activated as condensins dissociate and cells enter the next G1 phase. On the contrary, many chromosome-associated factors are known to be inherited through the cytoplasm (e.g., RNA polymerase, and CTCF) following mitotic exit. The evidence for contributions from both retained and re-acquired cis-regulatory programs to post-mitotic genome folding (reviewed in: (Ito and Zaret 2022; Budzyński et al. 2024) indicates that both modes of inheritance are likely occurring.

We have established an experimental system that allows us to discriminate the contributions of factors required for cell identity-specific chromosome folding based on whether they are inherited on mitotic chromosomes or transmitted to daughter nuclei through the cytoplasm. By preventing nuclear import from telophase through mitotic exit we were able to determine the extent to which interphase chromosome folding can be re-established without input from factors in the G1 cytoplasm. The folding patterns observed in the absence of nuclear transport must be determined and mediated by factors associated with chromatin during mitosis while any deviations from the expected conformation are caused by the inability of folding factors inherited through the cytoplasm to gain access to the genome after telophase. Our results demonstrate that the interphase chromatin state is achieved through the integration of two distinct folding programs inherited in distinct ways. We find that cell-identity specific instructions for compartmentalization of the genome are inherited on mitotic chromosomes and give rise to a folding intermediate driven by intrinsic affinities that is subsequently tuned by factors acquired from the cytoplasm, including cohesin, as cells exit mitosis.

## Results

### An experimental approach to insulate the genome from cytoplasmic factors during mitotic exit

During telophase of mitotic exit, the nuclear envelope re-forms around the segregated clusters of condensed chromosomes, and nuclear pore complexes (NPCs) are reassembled. Once the envelope has formed, cytoplasmic factors rely on regulated nuclear import across NPCs to gain access to the genome and associate with interphase chromosomes. We sought to prevent nuclear import from telophase onwards in order to understand the extent to which cell type-specific interphase chromosome folding can be re-established without input from factors in the G1 cytoplasm. To this end, two proteins essential for nucleocytoplasmic transport, RanGAP1 and Nup93, were biallelically fused with sequences for an auxin inducible degron (AID) and Neon Green fluorescent protein (NG) in DLD-1 cells using CRISPR/Cas9 gene editing (Methods). Nup93 is a scaffold nucleoporin essential for the assembly and stability of the nuclear pore complexes (Sachdev et al. 2012; Vollmer and Antonin 2014). RanGAP1 activates the Ras-related nuclear protein Ran in the cytoplasm and thereby critically contributes to the RanGTP gradient required for the nucleocytoplasmic shuttling of proteins (Quimby and Dasso 2003; Bischoff et al. 1994). These cell lines allowed us to inducibly target and disrupt nuclear transport by two different mechanisms.

In order to specifically prevent nuclear transport during mitotic exit, cells were synchronized and arrested in prometaphase using a double thymidine-nocodazole protocol (Supplemental Methods; Fig. 1a) and either RanGAP1-AID or Nup93-AID was efficiently degraded upon addition of auxin for 2 hours while cells remained arrested (Fig. 1b, Extended Data Fig.1b). Cells were then released from the nocodazole block and allowed to progress through anaphase and into G1. Depletion of RanGAP1-AID or Nup93-AID did not impact the kinetics of mitotic exit compared to untreated controls as evidenced by the loss of histone H3 phosphorylation after anaphase and the timely segregation of sister chromosomes to newly divided cells (Fig. 1c-d, Extended Data Fig.1a-b).

**Fig.1:**
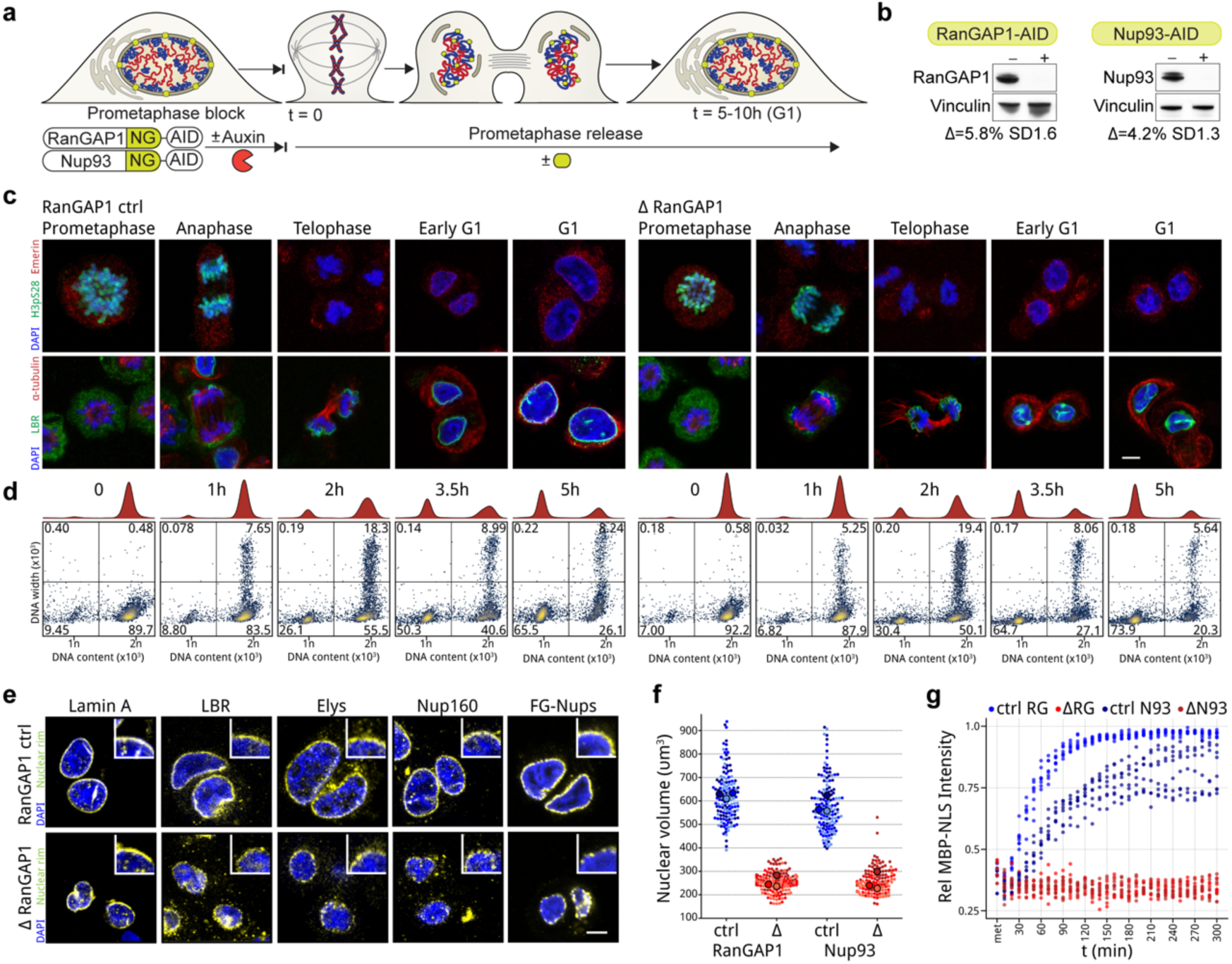
Acute depletion of essential nuclear import machinery during mitotic exit enables the assembly of daughter nuclei isolated from the G1 cytoplasm. a. Experimental workflow for depletion of either RanGAP1 or Nup93 AID-tagged proteins from the onset of mitotic exit into G1. Auxin-induced degradation is initiated 2h prior to nocodazole/mitotic release (t=0). b. Representative western blot images of whole cell lysates derived from control and auxin-treated DLD-1 cell lines 5 hours after mitotic release showing efficient depletion of RanGAP1-AID or Nup93-AID. Mean protein levels normalized to vinculin for depleted lysates relative to untreated controls are shown for 3 replicates with standard deviation (SD). c. Representative immunofluorescence images for RanGAP1 control and depleted cells demonstrate congruous kinetics of mitotic release. Loss of Histone H3 serine 10 phosphorylation (H3pS10) and recruitment of nuclear envelope proteins emerin and lamin B receptor (LBR) are shown for control and RanGAP1-AID-depleted cells. Scale bar represents 5 um. d. Representative DNA-content flow cytometry measurements for RanGAP1 control and depleted cells demonstrate congruous kinetics of mitotic release. e. Immunofluorescence images of RanGAP1-AID control and depleted cells 5 hours after mitotic release demonstrating presence of nuclear lamina (lamin A), nuclear envelope (LBR), and nuclear pore complex (Elys, Nup160, and FG-rich nucleoporins) resident proteins in the absence of RanGAP1-AID. Scale bar represents 5 um and inset indicates a 5x magnification of the nuclear rim. f. Nuclear volume measurements in control and RanGAP1 or Nup93-depleted G1 cells 5h after mitotic exit indicate the consistently smaller size of import-incompetent nuclei. Volume was calculated for 50 DAPI-stained nuclei per condition in three independent experiments. Individual data points and mean values for each replicate are indicated. g. Import competence is not re-established after mitosis in RanGAP1-AID or Nup93-AID-depleted cells indicated by the absence of nuclear import substrate accumulation. Relative nuclear to cytoplasm intensity of MBP-mScarlet-NLS is shown for eight cells in three independent fields taken every 10 minutes from the onset of mitotic release from Ro-3306 for one representative experiment. Data are plotted starting from metaphase, which is the first point on the graph, before cells proceeded to anaphase.

**Fig.2:**
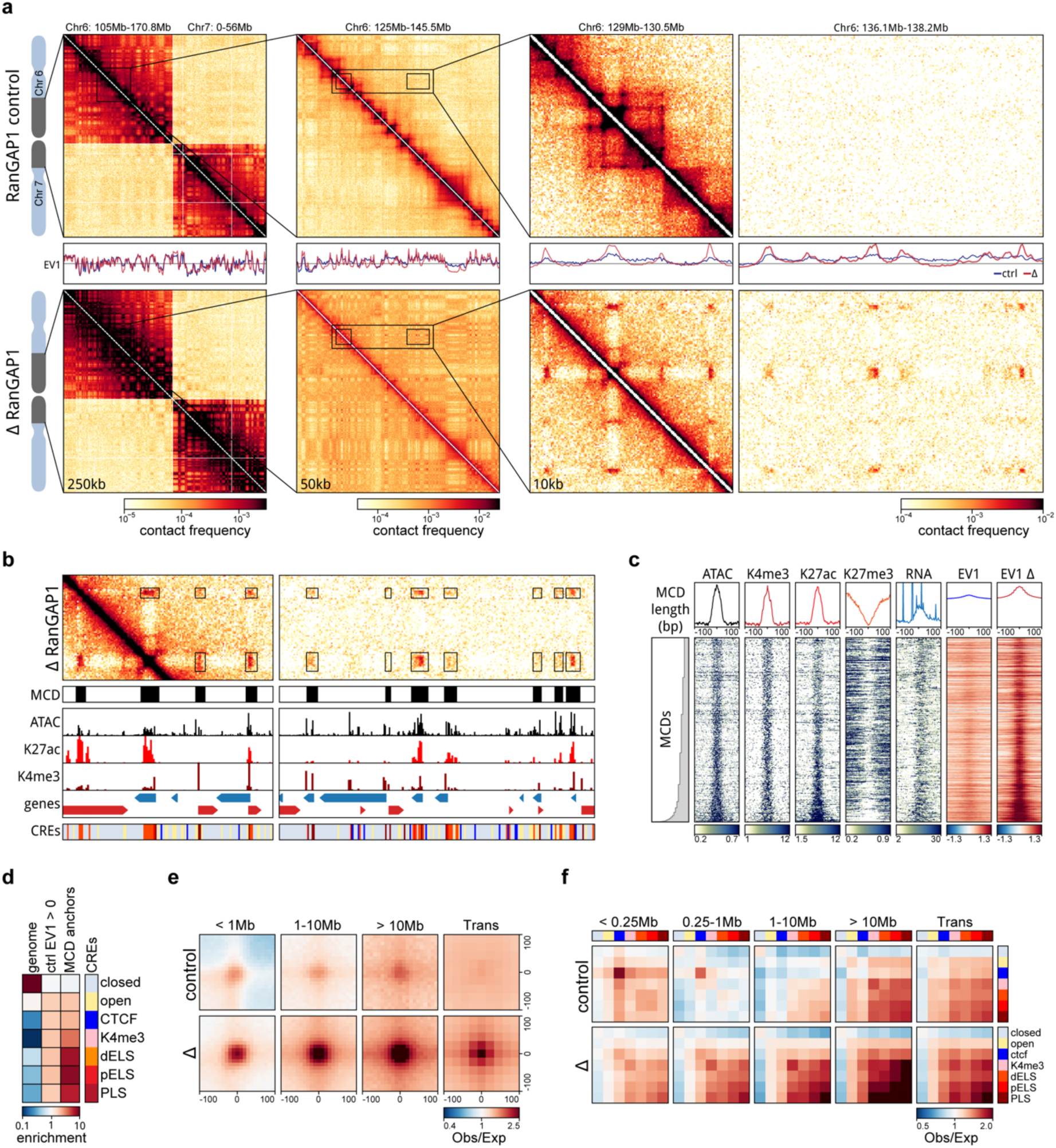
Depletion of RanGAP1 during mitotic exit reveals a chromosome-intrinsic capacity to form a kilobase scale cCRE compartment. a. Hi-C interaction frequency maps at 250kb (Chr 6: 105-170.8 Mb - Chr 7: 0-56 Mb), 50kb (Chr 6: 125 - 145.5 Mb), and 10kb (Chr 6: 129 - 130.5 & 136.1 -138.2 Mb) resolution showing genome compartmentalisation in control and auxin-treated DLD-1 RanGAP1-AID cells released from prometaphase arrest for 5h. Matched first eigenvector (EV1) values for cis interactions are phased by gene density (A > 0). a. b. Representative Hi-C contact matrix at 10kb resolution highlighting microcompartments in RanGAP1 depleted cells. Gene annotations, ATACseq, and histone H3 modification Cut&Run from control DLD-1 cells were used to define cis-regulatory elements (CREs), which coincide with microcompartment domains (MCDs). b. Heatmaps for all 2,105 MCDs centred on contact frequency summits and sorted by anchor length demonstrating the prevalence of control cell ATACseq, H3K27ac, H3K4me3 and H3K27ac coverage. Intra-chromosomal EV1 values from 10kb matrices indicate differences in compartmentalisation at MCDs in control and RanGAP1-depleted cells. c. Relative fold enrichment of control candidate cis-regulatory elements genome-wide, at 10kb control A-compartment bins (EV1 > 0), and at MCDs demonstrating the predominance of active promoters and enhancers at microcompartments. d. Pairwise mean observed/expected contact frequency between all MCDs projected in *cis* and *trans* showing dramatically enhanced interactions at all length scales and between chromosomes in RanGAP1-depleted G1 cells. e. Pairwise aggregate observed/expected contact frequency between cCREs assigned in control cells and subjected to hierarchical binning at 10kb resolution showing enhanced homo- and hetero-typic interactions between active promoters and enhancers in RanGAP1-depleted cells compared to controls at multiple genomic distances in *cis* and in *trans*.

The absence of Nup93 during the 5 hour mitotic release results in daughter nuclei sealed by a continuous nuclear envelope and underlying nuclear lamina that is devoid of nuclear pore complexes (Extended Data Fig. 1d-e), in accordance with an essential role for Nup93 in NPC assembly during mitotic exit. Conversely, apparently mature nuclear pore complexes that include the early recruited nucleoporins Elys and Nup160 as well as later associating FG nucleoporins (Mab414) are found at the nuclear envelope in RanGAP1-AID-depleted cells (Fig. 1e, Extended Data Fig. 1d). Despite these differences in nuclear morphology, depletion of either RanGAP1-AID or Nup93-AID impedes the extent of chromatin decondensation and nuclear volume expansion characteristic of control cells progressing through G1 (Fig. 1f, Extended Data Fig. 1d), suggesting that in both cases the interphase chromosome folding state is not fully realized. Furthermore, nucleoli and nuclear speckles, two import-dependent organelles with important roles in nuclear organization (Prasanth et al. 2003; Spector and Lamond 2011; Leung et al. 2004; Görlich and Kutay 1999), are not found in RanGAP1-AID or Nup93-AID-depleted nuclei and apparent interchromatin granules with strong speckle protein (SON) localization are found exclusively in the cytoplasm (Extended Data Fig. 1f).

In order to quantify the transport competence of large molecules in nuclei formed in the absence of Nup93 or RanGAP1, we repeated the depletion experiments in cells expressing a canonical importin-⍺ Nuclear Localization Signal (NLS) fused to Maltose-Binding Protein (MBP) and mScarlet fluorescent protein (MBP-mScarlet-NLS) (Fig. 1g, Extended Data Fig. 1f). Time lapse fluorescent imaging of control G2-synchronized cells released into mitosis demonstrates efficient nuclear import of MBP-mScarlet-NLS following chromosome segregation, with minimal protein observed in the cytoplasm by 5 hours when cells are in G1. In contrast, depletion of either Nup93-AID or RanGAP1-AID entirely prevents nuclear accumulation of MBP-mScarlet-NLS throughout mitotic exit and G1 entry, consistent with a failure to establish nucleocytoplasmic transport. Thus, in this experimental context, a nucleus is re-built during mitotic exit around newly segregated decondensing chromosomes, but the genome is completely isolated from the cytoplasm once the nuclear envelope has formed in late telophase.

### Post-mitotic chromosomes possess the intrinsic capacity to form a genome-scale microcompartment of cCREs

To assess how chromosome-intrinsic factors inform post-mitotic genome folding, we performed Hi-C 3.0 (Lafontaine et al. 2021; Akgol Oksuz et al. 2021) with G1 cells that had progressed through mitosis in the absence of either RanGAP1 or Nup93 (Supplementary Table 1, Supplementary Fig. 1a). Newly divided cells were fixed 5 hours after nocodazole release and fluorescence-activated cell sorting (FACS) was employed to obtain pure G1 populations. Despite an inability to establish nucleocytoplasmic transport, A/B compartments are not only formed but appear visually more pronounced in RanGAP1-AID and Nup93-AID-depleted cells (Fig. 2a, Extended Data Fig. 2a; see below). Global compartmentalization, captured by the first Eigenvector (EV1) of the Hi-C contact matrix, is similar in nuclear import-deficient nuclei compared to control DLD-1 cells (Supplementary Fig. 1b), suggesting that the determinants and mediators of cell type-specific compartmentalization are already associated with chromosomes by telophase. In the absence of nuclear import, we observe an additional prominent network of small (< 100 kb) highly interacting chromatin domains that spans entire chromosomes. These domains interact ubiquitously with each other in both cis and in trans. The grid-like nature of these interactions implies the formation of a chromosome-intrinsic compartment. These interactions form independently of factors that would normally enter the nucleus after telophase.

We explored the properties of this chromosome-intrinsic microcompartment by identifying the domains of 25-125 kb that give rise to strong focal pairwise interaction enrichments in the Hi-C interaction data of RanGAP1-AID depleted G1 cells. Custom modification of the convolutional kernels and pixel filtering steps previously used to call loops in Hi-C data ((Rao et al. 2014; Open2c et al. 2024), Supplementary Fig.2, Supplemental Methods) identified 2,105 strong microcompartment domains (MCDs) encompassing all autosomal chromosomes. Microcompartment domains encompass gene dense active regions corresponding to high ATACseq coverage as well as peaks of histone H3K4-trimethylation and H3K27-acetylation (Supplementary tables 2-3) found in unperturbed DLD-1 cells (Fig. 2b-c). Accordingly, active promoters and enhancers defined in control cells (candidate cis-regulatory elements: cCREs, see methods) are highly enriched at MCDs even when comparing their relative enrichment in the active compartment (“EV1 > 1”) (Fig. 2d). The first Eigenvector indeed indicates an A-compartment profile at MCDs in control cell lines. In RanGAP1-AID or Nup93-AID-depleted cells EV1 becomes sharper with prominent peaks aligned to MCDs (Fig. 2a,c; Extended Data Fig. 2a,b) indicating the emergence of a finer-scale microcompartment. We refer to MCDs as containing active elements because of the prominence of active chromatin marks and active promoters defined in control cells. However, we note, and show below, that much of the transcriptional machinery is absent from the nucleus in import-deficient cells and thus the prominent compartmentalization of these active elements from the previous cell cycle likely occurs in the absence of ongoing transcription.

Based on the comprehensive grid of microcompartment domain interactions observed in RanGAP1-AID-depleted cells, we assumed that all MCDs possess an intrinsic affinity for each other. In order to quantify their interactions, we projected all pairwise combinations of MCDs to give rise to 73,970 and 2,079,615 pairwise microcompartment interactions within and between chromosomes, respectively (Supplemental methods). Aggregate pile-ups demonstrate at least a 4-fold average focal enrichment in interaction frequency at projected pairs of MCDs in RanGAP1-AID or Nup93-AID depleted cells extending to all genomic length scales in *cis* and to *trans*-chromosomal contacts (Fig. 2e, Extended Data Fig. 2c). The quantitatively lower enrichment of MCD contact frequency in Nup93-AID-depleted cells compared to RanGAP1 depletion may reflect clonal variation between the two cell lines as MCDs were assigned in the latter context, but we can not exclude the possibility that blocking nuclear pore complex assembly confers additional consequences for post-mitotic nuclear organization beyond the effect on nuclear import. Nonetheless, these data suggest that clustering of MCDs is a general and prominent feature of post-mitotic chromosome folding in the absence of nuclear import.

Enriched interactions between the MCDs detected in Ran-GAP1-AID depleted cells are also observed in control G1 cells but with substantially reduced frequency (Fig. 2e). The capacity for MCDs to interact in wildtype G1 cells is consistent with previous observations of weakly enriched interactions between active promoters and enhancers (Friman et al. 2023; Harris et al. 2023; Goel et al. 2023).

In a complementary analysis we determined interaction frequencies between cCREs defined in wildtype DLD-1 cells at 10 kb resolution. As expected, pairs of CTCF sites interact frequently in control cells at relatively small genomic distances (< 1 Mb). In contrast, CTCF-CTCF interactions are not enriched in RanGAP1-AID or Nup93-AID depleted G1 cells (Fig. 2f), addressed in more detail below. We found that, like MCDs, active promoters and enhancers interact with each other at considerably elevated frequencies regardless of genomic separation when nuclei are formed in the absence of RanGAP1 or Nup93 (Fig. 2f, Extended Data Fig. 2d). In control cells these interactions also occur, but at much lower frequencies. Taken together, we find that the propensity to form a microcompartment of active cis-regulatory elements during G1 is an intrinsic capacity of post-mitotic chromosomes that emerges as a much more pronounced feature in the absence of nuclear transport.

### A microcompartment of cis-regulatory elements is a transient folding intermediate during mitotic exit

Telophase is the latest stage of mitotic exit when chromosomes can recruit cytoplasmic factors without a requirement for nuclear import and is the last point of common chromosome composition between control and RanGAP1-AID or Nup93-AID-depleted cells entering G1. We therefore sought to determine the folding state of telophase chromosomes using Hi-C. The brevity of telophase and lack of a robust exploitable check-point poses a particular obstacle in obtaining synchronous populations of cells in this phase. To address this challenge, we first arrested cells in prometaphase and released them from the nocodazole block for 1.25 or 1.5 hours, as we had previously observed the greatest proportion of telophase cells in the post-mitotic population between 1 and 2 hours (shown in Fig. 1d, upper right quadrant).

To enrich telophase populations of control and RanGAP1-AID-depleted cells, fixed cells were stained with propidium iodide (PI) and sorted based on the intensity and the width of the PI signal (Fig. 3a), which briefly peaks in the transition from metaphase to G1 ((Gasnereau et al. 2007), also seen in Fig. 1c). Manual scoring of immunofluorescent images for the isolated cell populations indicated successful enrichment of two distinct cell cycle stages during this critical mitotic exit window. At 1.25 hours post release, at least 70% of sorted cells were in telophase, as evidenced by elongated midbody microtubules connecting the two masses of highly condensed chromosomes. Cells sorted in the same way at 1.5 hours post release had progressed through telophase and we could identify close to 60% of cells having a clear abscission midbody, indicating they had not yet completed cytokinesis. In our experimental system, nuclear volume increases by about 50% from telophase to cytokinesis in control cells and dramatically expands by close to 3-fold in early G1 (Fig. 3b). In contrast, RanGAP1-AID-depleted nuclei do not noticeably expand in size beyond telophase.

**Fig. 3:**
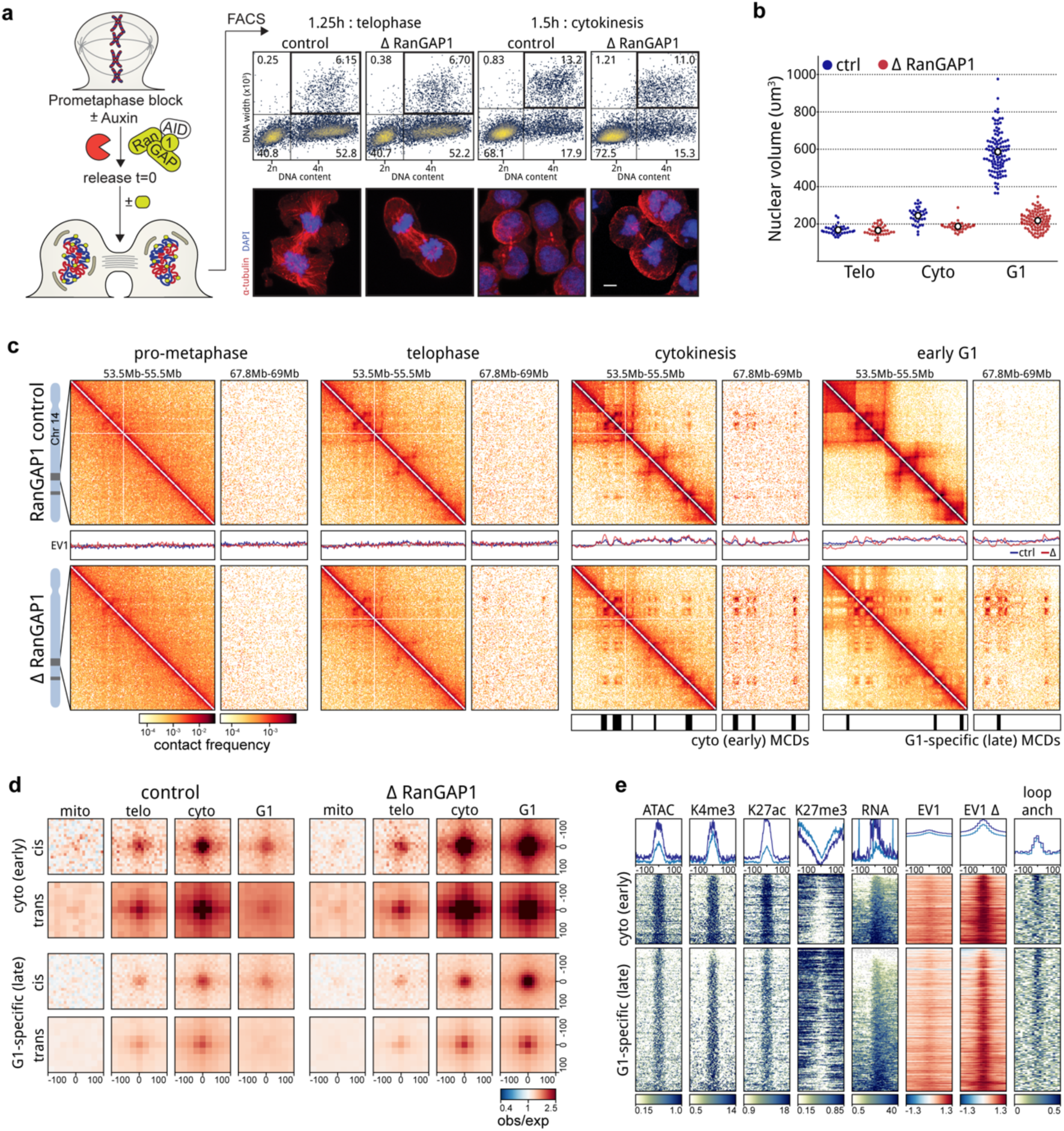
A transient microcompartment of chromosome-intrinsic affinities is first formed during telophase. a. Isolation of highly synchronous telophase and cytokinesis cell populations by FACS for Hi-C analysis. Fixed cells are collected 1.25-1.5h following prometaphase release and sorted based on DNA (PI) content and width. Auxin-induced degradation of RanGAP1 is initiated 2h prior to nocodazole release (t=0). Representative immunofluorescence images show the enrichment of cells in telophase or cytokinesis based on the morphology of chromatin (DAPI) and microtubules (alpha-tubulin). Scale bar represents 5 um b. Nuclear volume measurements in control and RanGAP1-depleted cell populations enriched in telophase, cytokinesis, or early G1 indicate a lack of nuclear growth after telophase in import-incompetent nuclei. Volume was calculated for 25 DAPI-stained nuclei per condition. c. Representative Hi-C interaction frequency maps at 10kb (Chr 14: 53.5 - 55.5 v. 67.8 - 69 Mb) resolution showing genome organisation in control and auxin-treated DLD-1 RanGAP1-AID cells enriched in prometaphase, telophase, cytokinesis, or early G1 (5h after release). Matched first eigenvector (EV1) values for cis interactions are phased by gene density (A > 0). d. Pairwise mean observed/expected contact frequency between MCD anchors showing enhanced interactions in telophase that peak around cytokinesis of mitotic exit in control cells. All 2,105 G1 MCD anchors are categorized by whether they are detected early (cytokinesis) or late (G1-specific) in RanGAP1-depleted cells and projected in *cis* and *trans*. e. Heatmaps for MCD anchors, divided by whether they are detected early (cytokinesis) or late (G1) in RanGAP1-depleted cells. Stacks centred on contact frequency summits and sorted by MCD anchor length demonstrate the prevalence of control cell ATACseq, H3K27ac, H3K4me3 and H3K27me3 coverage and differences in intra-chromosomal in 10kb EV1 values for control and RanGAP1-depleted G1 cells at early and late MCD anchors. Plots on top of the heatmaps represent average profiles of the corresponding features, with the dark blue line representing the MCDs detected in cytokinesis and the light blue lines representing the G1 MCDs.

We performed Hi-C with FACS-sorted populations of prometaphase, telophase, cytokinesis, and G1 cells and observed the progressive acquisition of interphase chromosome compartments and loops in control cells (Fig. 3c). Remarkably, unperturbed cells were found to form microcompartment domain contacts that are visually apparent in contact frequency heatmaps by cytokinesis. We classified MCDs based on whether they were detected early (cytokinesis) or late (G1-specific) during mitotic exit in the RanGAP1-depleted condition. Aggregate analysis demonstrates that pairwise interactions between these early and late MCDs are formed in *cis* and *trans* during telophase and increase in strength towards cytokinesis in both control and RanGAP1-AID-depleted cells (Fig. 3d). In the absence of nuclear import, this active microcompartment continues to strengthen as cells enter G1. By contrast, progression into G1 in control cells coincides with a strong reduction in the frequency of MCD contacts. Together, these data show that extensive *cis* and *trans* micro compartmentalization between active cCREs is a normal transient process that peaks during cytokinesis and is then lost or at least severely reduced. In RanGAP1-AID or Nup93-AID-depleted cells this loss is not observed and MCD interactions continue to increase in frequency as cells progress through G1, suggesting that import of factors from the cytoplasm normally overrules or counteracts this process The specific set of MCDs identified in RanGAP1-depleted cells specifically at cytokinesis are not unique to this cell cycle stage but rather represent a subset of all G1 MCDs that interact with greater contact frequency throughout the time course (Fig. 3d). These early detected MCDs are relatively more enriched for marks of

active cis-regulatory elements compared to the MCDs detected later in G1 (Fig. 3e, Extended Data Fig. 3a), consistent with the formation of an active cCRE-driven microcompartment during mitotic exit. Accordingly, contacts between active promoters and enhancers mirror interactions between MCDs during mitotic exit (Extended Data Fig. 3b). Average contact frequency between cis-regulatory elements increases in telophase reaching a peak at all length scales during cytokinesis and is relatively diminished in a distance-dependent way in control cells by early G1.

### Microcompartment domains are enriched for mitotically bookmarked gene regulatory elements

Cell identity is propagated to daughter cells by epigenetic chromatin modifications and retained protein factors that mark active promoters and some enhancers during mitosis (Ito and Zaret 2022). The appearance of a microcompartment during telophase that overlaps the specific active cis-regulatory elements of wild type asynchronous cells implies an inherited feature. To investigate this possibility, we performed omni-ATACseq (Grandi et al. 2022) for cells arrested in prometaphase or released for 5 hours to early G1 in the presence or absence of RanGAP1 (Supplementary table 2). In agreement with previous work (Oomen et al. 2019), ATACseq fragment length distributions indicate more regularly spaced nucleosomes during mitosis and a presumably more dynamic, less regularly arranged nucleosome landscape in G1 (Extended Data Fig. 4a). Depletion of RanGAP1-AID did not affect the difference in fragment length distribution between mitotic and G1 cells, despite differences in chromosome decondensation (Fig. 1f).

We identified 159,072 peaks of ATACseq coverage across all conditions and compared their intersection (Fig. 4a). The majority of peaks found genome-wide are specific to G1 cells, consistent with a general loss of accessibility during mitosis that is regained during G1. The largest fraction of open regions are G1-specific peaks that open during mitotic exit in control cells, but that do not become accessible in RanGAP1-AID-depleted cells entering G1 (40%). This suggests that the opening of G1-specific sites generally requires factors that are inherited through the cytoplasm and only gain access to chromatin after nuclear envelope formation in G1. The second largest fraction of open regions are mitotically retained peaks that are common to all conditions (30%) and represent constitutively open (bookmarked) sites.

**Fig. 4:**
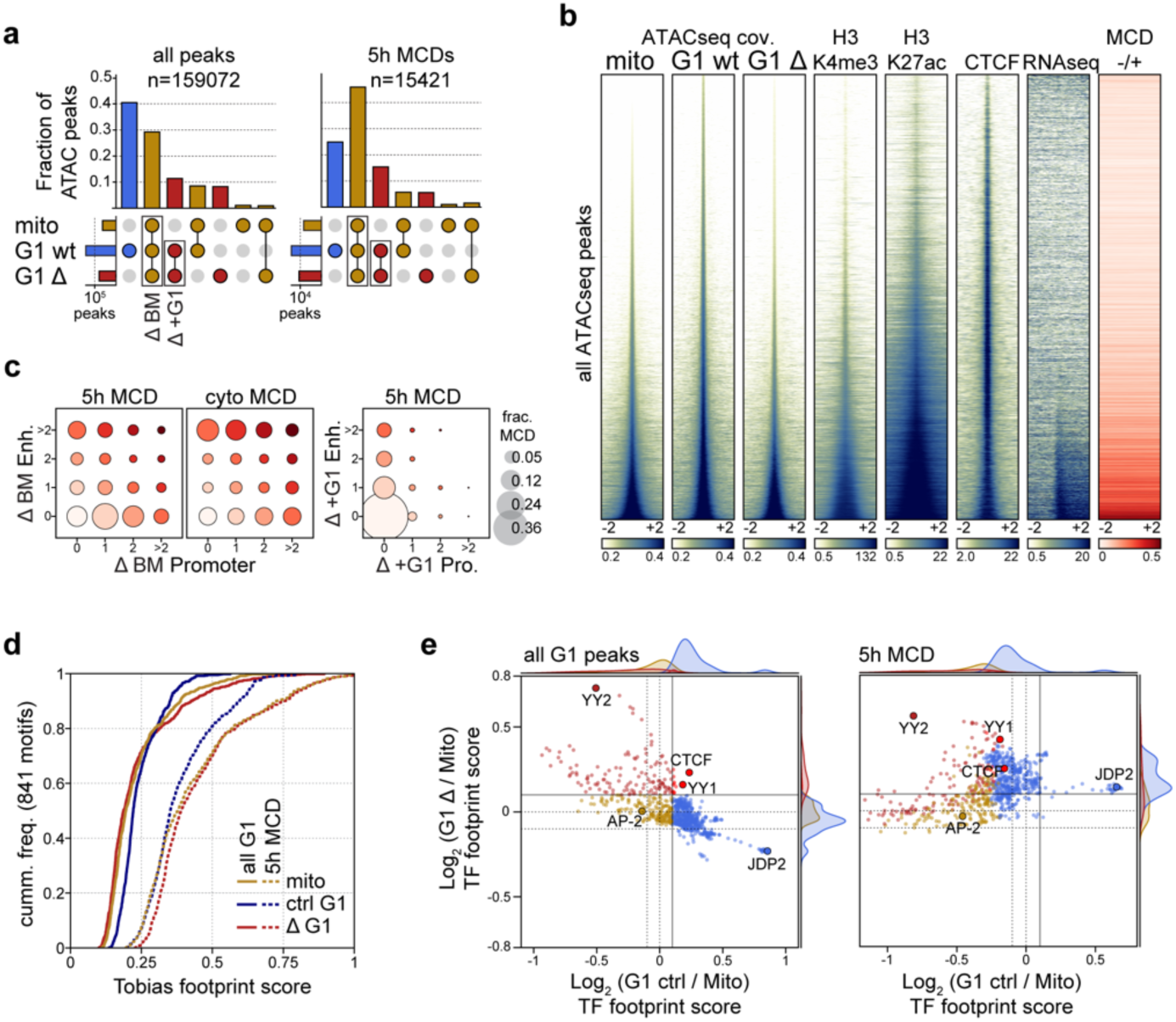
Microcompartment domains are bookmarked during mitosis. a. Intersections of all (left) or MCD-overlapped (right) ATACseq peaks detected in prometaphase, G1 control, or RanGAP1-depleted G1 cells are plotted as a fraction of the total. The number of called peaks are indicated. Bookmarked (BM) and G1-specific peaks present in RanGAP1-depleted cells are highlighted. b. Heat Maps centred on the union set of ATACseq peaks detected in prometaphase-arrested or G1 cells released for 5h -/+ RanGAP1. Stacks sorted by prometaphase ATACseq signal demonstrates co-occurence of G1 signals at mitotic peaks and prevalence of H3K27ac, H3K4me3, RNAseq, and MCD anchors. c. The proportion of all G1 MCD anchors or the cytokinesis subset at the indicated numbers promoters and enhancers demonstrates the valency of bookmarked (BM) and G1-specific CREs. Bubble size indicates the relative frequency. d. Empirical cumulative density plot of TOBIAS TF footprinting scores for 841 non-redundant vertebrate Jaspar motifs calculated for prometaphase or G1 -/+ RanGAP1 ATACseq signal at all control G1, or MCD-overlapped peaks. e. Comparison of differential TOBIAS footprint scores for either control or RanGAP1-depleted G1 vs prometaphase ATACseq at all G1 or MCD-overlapped G1 peaks. Each dot represents one of 841 non-redundant vertebrate Jaspar motifs, coloured by the distribution of scores at all G1 peaks. Bookmarked (gold) and G1-specific (blue) motifs are further classified by higher footprinting scores in the RanGAP1 depletion (dark and bright red, respectively) and relative density is plotted for each category for the control (x-axis) and depletion (y-axis) Log2-fold change over prometaphase.

The composition of ATAC-seq peaks overlapping MCDs differs substantially from the genome-wide peaks. At MCDs, accessible regions are predominantly bookmarked peaks (i.e., also accessible in mitosis), while G1-specific peaks are relatively infrequent. Retained mitotic peaks correspond to highly accessible chromatin at all cell cycle stages and coincide with the promoters of active genes and strong H3K27ac signal in control DLD-1 cells (Fig. 4b, Extended Data Fig. 4b-c), consistent with mitotic bookmarking at cis-regulatory elements. More than 90% of MCDs overlap at least one bookmarked promoter- or enhancer-annotated ATACseq peak and the valency of these transcriptional elements (i.e., the number of MCD interactions it is engaged in RanGap1-AID-depleted cells) is increased at the stronger domains that were detected earlier, in cytokinesis (Fig. 4c). A small subset of G1-specific peaks that become accessible in the import-deficient G1 cells are also enriched at MCDs, implying a potential contribution from elements acquired after mitotic exit to MCD formation. However, more than half of the MCDs do not overlap any promoter or enhancer specific to these G1-specific peaks, and thus these G1-acquired open sites are not essential for MCD interactions. Taken together, these data suggest that the microcompartments formed during telophase are likely to be driven by mitotically inherited regions of robust chromatin accessibility.

An increasing number of transcription factors have been proposed to act as cell type-specific bookmarks of gene regulatory information during mitosis (Palozola et al. 2019). Our ATACseq data confirms the prevalence of accessible active promoters during mitosis and indicates a high density of bookmarked promoters and enhancers at the microcompartment domains detected in the RanGAP1-AID depletion. In an effort to characterize the cell cycle dynamics of this epigenetic landscape, transcription factor (TF) footprinting was investigated at 841 conserved vertebrate motifs (Fornes et al. 2020) using the orthogonal ATACseq analysis approach TOBIAS (Bentsen et al. 2020). Genome-wide footprint scores at most TF motifs are stronger in the control G1 condition compared to mitosis, affirming that most sites are closed during mitosis and acquire accessibility in G1 (Fig. 4a). Footprint scores within MCDs are considerably higher than at all open regions genome-wide, both in G1 controls and in mitosis (Fig. 4d). This observation is consistent with MCDs containing bookmarked open sites. In RanGAP1-AID depleted cells, we did not observe broad cell cycle-dependent dynamics in footprinting. Average footprint scores across TF motifs in RanGAP1-depleted G1 cells were largely unchanged compared to mitosis, with MCDs containing the strongest footprints. This analysis confirms that wildtype cells display genome-wide dynamics in TF binding site accessibility as cells enter G1, while the accessibility landscape of TF binding sites in nuclear import deficient G1 cells reflects the bookmarked mitotic state.

To determine whether specific transcription factors contribute to bookmarking at MCDs, we compared the change in accessibility of each individual motif as control or RanGAP1-depleted cells exit mitosis. As expected, the majority of motifs genome-wide have increased average footprinting scores in control G1 cells compared to mitosis while very few motifs become more accessible during G1 in the absence of RanGAP1 (Fig. 4e left panel). The small group of motifs with increased average footprints in RanGAP1-AID-depleted cells includes two proteins of particular interest to chromosome folding: CTCF and YY1, for which footprints are also increased in control G1 cells (Extended Data Fig. 4d). This is consistent with the observation that CTCF binding occurs rapidly after mitotic exit (around telophase) and prior to nuclear envelope closure. We also note that the YY2 motif gains footprinting strength specifically in RanGAP1-AID-depleted cells. Cell cycle-dependent differences in footprinting at TF motifs specifically within MCDs are less pronounced compared to those found genome-wide (Fig. 4e right panel), consistent with the increased accessibility of bookmarked sites during mitosis. Notably, as footprint scores are normalized across all control G1 peaks, the distribution of differential control G1 footprints is globally shifted towards stronger mitotic footprinting at MCDs, where a high concentration of these bookmarked sites are found. At MCDs, the relative changes in footprinting scores in RanGAP1-depleted cells are better correlated with control footprints, suggesting that opening or binding of the TF motifs within these domains is not dependent on nuclear transport. Taken together, this analysis demonstrates that most TF motifs genome-wide gain footprinting strength as cells enter G1, and this gain is dependent on nuclear import. Conversely, TF binding sites found at MCDs demonstrate relatively strong footprints even in mitosis that do not appear to strengthen during G1. Specific TF motifs do not emerge at MCDs but rather there is a convergence to strong footprinting in all conditions regardless of whether a particular motif is robustly bookmarked (i.e., TFAP2) or G1-specific genome-wide (i.e. JDP2) (Extended data Fig. 4d-e), suggesting that the architecture in these domains already reflects the G1 epigenetic state throughout the cell cycle.

### Nuclear transport-dependent pruning of long-range indiscriminate microcompartment interactions

At the end of mitosis, around telophase (Zhang et al. 2019; Abramo et al. 2019; Brunner et al. 2024), the condensin-mediated loop array that compacts mitotic chromosomes is replaced by the extrusive activity of cohesin, which gives rise to CTCF-CTCF loops, TADs, and insulating domain boundaries. Previous Hi-C studies have demonstrated the progressive acquisition of these features during G1 entry, concomitant with the accumulation of cohesin on the chromatin (Zhang et al. 2019; Abramo et al. 2019). The average contact probability between pairs of loci as a function of their genomic separation, *P*(*s*), can be used to estimate the size of extruded loops. The shape of *P*(*s*) typically displays a “bump” where interaction frequency decays the slowest. This distance of minimal decay in contact frequency represents the average loop size (Gassler et al. 2017; Gibcus et al. 2018; Schwarzer et al. 2017; Polovnikov et al. 2022; Haarhuis et al. 2017) and can be readily identified as a local peak in the derivative *P*(*s*) plot. In prometaphase, the average loop size is around 200-300 Kb, regardless of RanGAP1-AID depletion. These ∼200-300 kb condensin loops disappear at telophase in control DLD-1 cells and are mostly undetectable by cytokinesis. As control cells enter G1, loops with an average size of ∼100 kb appear, as expected for loops generated by cohesin (Fig. 5a).

**Fig. 5:**
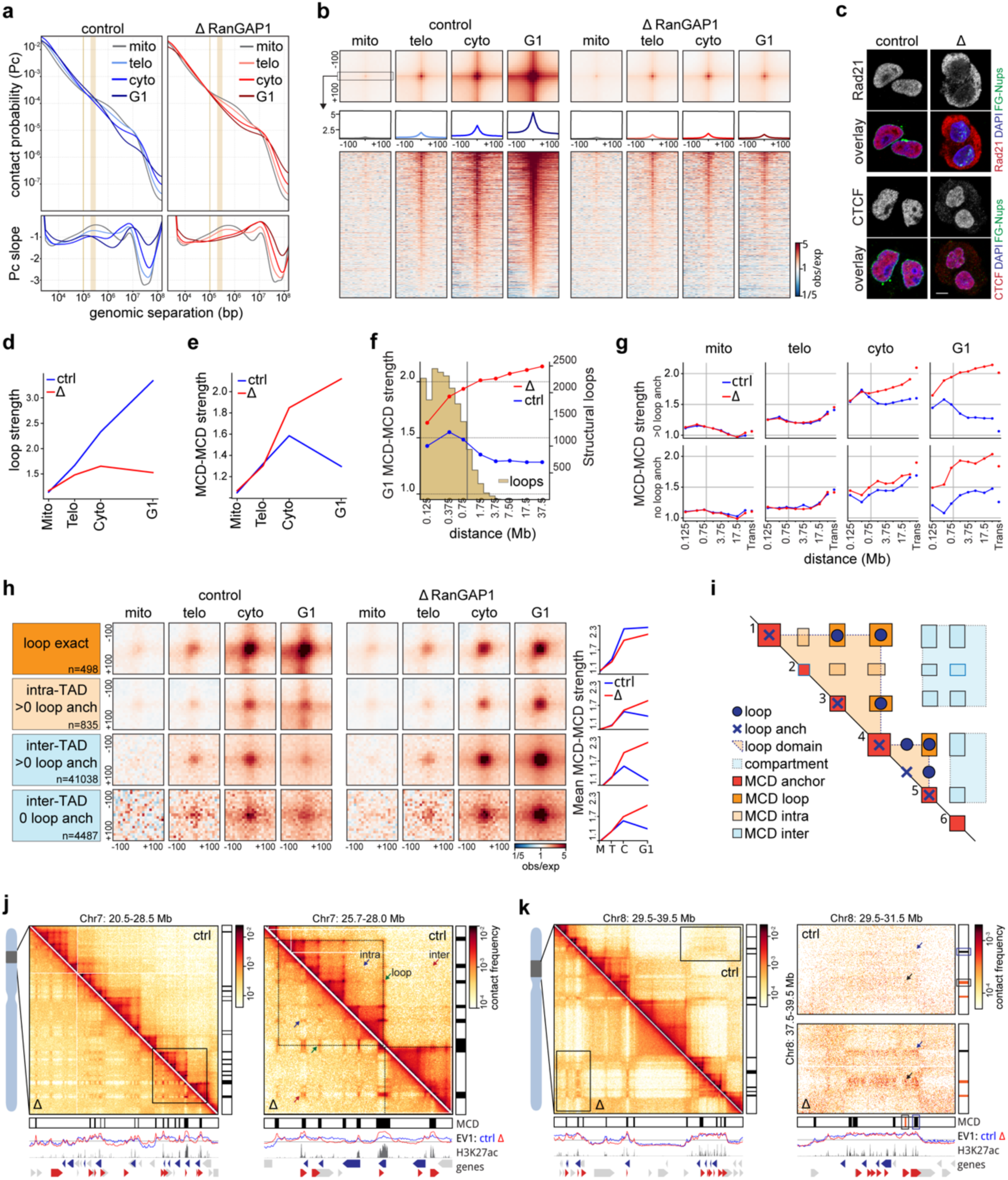
Microcompartment interactions are pruned in nuclear transport-competent cells by early G1. a. P(s) and derivative P(s) plots for Hi-C data from synchronised control and RanGAP1-depleted FACS-sorted prometaphase, telophase, cytokinesis, and early G1 (t = 5h) cells. b. Mean observed/expected Hi-C contact frequency at 18,613 convergent CTCF loops identified in pooled interphase G1 control Hi-C data demonstrating the RanGAP1-dependent increase in looping interactions as cells progress from prometaphase to G1. Average signal for three central 10kb bins across the 200kb CTCF motif-centered window and stack-ups sorted by G1 loop strength are shown. c. Representative immunofluorescence images of RanGAP1-AID control and depleted cells fixed 5h after mitotic release demonstrating the nucleo-cytoplasmic localisation of the cohesin complex subunit, Rad21, and boundary transcription factor, CTCF. Scale bar represents 5 um. d. Mean convergent CTCF loop strength quantified for RanGAP1-AID control and depleted cells (panel e) at discrete points of mitotic exit, as indicated. e. Mean strength of all pairwise MCD-MCD contacts projected in *cis* and quantified for RanGAP1-AID control and depleted cells in early G1 (t = 5h) at different distances of separation demonstrating the distance-dependent reduction in interaction strength beyond the range of structural CTCF loops. f. Mean strength of MCD-MCD contacts projected in *cis* and quantified for RanGAP1-AID control and depleted cells at discrete points of mitotic exit and across separation distances. Pairwise MCD interactions are classified based on the presence (l.a. > 0) or absence (l.a. = 0) of at least one CTCF loop anchor. g. Pairwise mean observed/expected contact frequency between MCDs at discrete points of mitotic exit, as indicated, in control and RanGAP1-depleted cells. MCD-MCD contacts are categorised by the presence of a convergent CTCF loop or loop anchor (> 0) and the looping domain status of the constituent MCDs, as indicated. h. Schematic summary of microcompartment fates in G1 cells based on the spatial relationships of constituent MCDs to extrusion-dependent features. i. Representative Hi-C interaction frequency maps at 25kb (Chr7: 20.5-28.5 Mb) resolution and a 10kb zoom-in (Chr7: 25.7-28.0 Mb),showing differences in genome organisation between two looping domains in control (upper triangle) and RanGAP1-AID-depleted G1 cells (t = 5h, lower triangle). Matched MCD position tracks, as well as control cell EV1 values, H3K27ac Cut&Run signal, and gene annotations are shown. j. Representative Hi-C interaction frequency maps at 50kb (Chr8: 29.5-39.5 Mb) and 10kb (Chr8: 29.5-31.5 Mb v. 37.3-39.5 Mb) resolution showing differences in genome organization at loop anchored and extrusion-free MCDs in control and RanGAP1-AID-depleted G1 cells (t = 5h). Matched MCD position tracks, as well as control cell EV1 values, H3K27ac Cut&Run signal, and gene annotations are shown.

When RanGAP1-AID is acutely depleted during mitotic exit, condensin loops are lost with similar kinetics to the control but ∼100 kb loops are not established and the *P*(*s*) curve indicates the absence of any extrusion loops in cytokinesis and G1. These observations are consistent with the existence of a transient folding intermediate during mitotic exit that lacks both condensin- and cohesin-mediated loops (Abramo et al. 2019). We conclude that the appearance of interphase extruded loops, mediated by cohesin, requires nuclear import and thus initiates after nuclear envelope formation. As cohesin-extruded loops are frequently anchored at sites bound by CTCF in an orientation-dependent way (Fudenberg et al. 2016; Sanborn et al. 2015; Rao et al. 2014), we identified the set of all control interphase loops at convergent CTCF motifs and quantified the interactions between these loci during mitotic exit. The strongest interphase loops first become detectable, albeit weakly, at telophase in both control and RanGAP1-depleted cells. However, while the mean looping strength increases considerably during cytokinesis and reaches at least 5-fold enrichment in early G1 control cells, loops remain rare and very weak throughout mitotic exit in the absence of RanGAP1 (Fig. 5b-d). The weak loop signal observed in RanGAP1-AID-depleted telophase cells may be the result of a very small amount of cohesin loading onto chromatin before the nuclear envelope closes. Depletion of Nup93-AID during mitotic exit similarly prevents the establishment of interphase loops (Extended Data Fig. 5a-b), suggesting that functional nuclear import is a general requirement for cohesin-mediated loop extrusion during exit from mitosis and entry into G1. Consistently, immunofluorescence indicates the exclusion of the cohesin complex kleisin subunit, Rad21, from post-mitotic nuclei assembled in the absence of either RanGAP1 or Nup93, whilst CTCF retains at least partial access to chromatin in the nucleus (Fig. 5c, Extended Data Fig. 5c). Accumulation of CTCF on the chromatin in the absence of nuclear import is consistent with our ATACseq data and supports previous findings that suggest the rapid recruitment of CTCF at mitotic exit precedes the cohesin-mediated extrusion of interphase loops (Zhang et al. 2019). Our data demonstrates that CTCF re-binds chromatin prior to nuclear envelope closure.

Loop extrusion has been found to counteract epigenetic-defined chromatin compartmentalization (Schwarzer et al. 2017; Haarhuis et al. 2017; Nuebler et al. 2018; Rao et al. 2017). The cell cycle dynamics of the cCRE microcompartment in control cells adhere to this logic. We observe a decrease in MCD interaction strength in control cells concomitant with an increase in the strength of extruded loops as cells enter G1. Conversely, pairwise MCD interactions continue to increase in frequency as RanGAP1-depleted cells progress to G1 in the absence of extruded loops (Fig. 5d-e).

The reduced pairwise MCD interaction strength in control G1 cells is not uniform across genomic distances (Fig. 5f). Short range (< ∼500 kb) MCD-MCD contacts are preferentially retained, while pairwise interaction strength drops substantially for MCDs separated by larger distances, beyond the range of most extruded loops. These data suggest that the indiscriminate pairwise MCD interactions appearing from telophase to cytokinesis are specifically lost at large genomic distances in control G1 cells. We hypothesized that this phenomenon was directly related to cohesin-mediated loop extrusion. To address this supposition, microcompartment domains were categorized based on whether they overlap a CTCF loop anchor. At 10 kb resolution, extrusion anchors are found at more than 75% of MCDs. We compared the strength of pairwise MCD interactions (focal interaction frequency over local background, Supplemental Methods) that coincide with at least one extrusion anchor to the MCD contacts that do not have any CTCF loop anchor as a function of their genomic separation during mitotic exit (Fig. 5g). For both sets of MCDs, pairwise interaction strength increases from prometaphase to cytokinesis in control as well as in RanGAP1-AID-depleted cells (Fig. 5g). MCD-MCD interactions continue to strengthen into G1 in RanGAP1-AID-depleted cells for all genomic distances and regardless of loop anchor status. However, in control cells the two sets of MCDs diverge in their distance-dependent interaction frequencies from cytokinesis onwards. MCD interactions overlapping at least one CTCF loop anchor are maintained only when they are less than 500 kb apart, while contacts separated by larger distances become much less frequent in G1. The differential strength of long and short-range MCD interactions is already detectable at cytokinesis when extrusion starts but while short range interactions are almost unchanged as cells progress to G1, long range contacts forming microcompartments are further reduced. Conversely, MCD interactions that do not overlap any CTCF loop anchor do not demonstrate a distance dependent loss of interaction frequency in control cells and from cytokinesis to G1 focal contact frequencies are maintained at larger distances.

These findings imply a role for loop extrusion in modifying the global network of pairwise MCD-MCD interactions as cells exit mitosis, through a process we refer to as “pruning”. Pairwise MCD interactions that appear in cytokinesis and coincide with interphase CTCF loops are not only retained but also strengthened in control G1 cells (Fig. 5h-j, “loop exact”), suggesting that while initial MCD-MCD interactions are cohesin-independent, cohesin-mediated loop extrusion specifically reinforces these contacts. The fact that cohesin loops are typically less than 500 kb can explain why only relatively short-range microcompartments are maintained in this manner while longer-range contacts are reduced, or pruned, in G1 (Fig. 5f). As these MCDs become tethered to a small set of nearby MCDs to give rise to interphase extrusion loops, interactions with other more distal MCDs become far less probable.

To more specifically explore extrusion-dependent pruning, we further classified the remaining microcompartmental interactions based on the domain identity of constituent MCDs. An extrusion domain or TAD was defined in control G1 cells by considering the outermost loop from any subgroup of nested convergent CTCF loops (Supplemental Methods). Interactions between pairs of MCDs that overlap at least one CTCF loop anchor and that span two distinct domains (inter-TAD) are pruned as control cells progress from cytokinesis to G1. Conversely, interactions between pairs of MCDs that are both located within a single domain but not overlapping an extrusion loop (intra-TAD) are only slightly diminished (Fig. 5h-j). The intra-TAD MCD interactions may reflect weaker extrusion loops or nested loops that favor the outermost interaction in G1 or may become more diffuse because of extrusion-mediated mixing within the TAD. Inter-TAD interactions between pairs of MCDs that do not harbor any CTCF loop anchors are also reduced as control cells exit mitosis (Fig. 5h-i,k) but to a much lesser extent than those overlapping a CTCF loop anchor, suggesting that proximity to the extrusion anchor may have a greater influence on pruning than the domain architecture. In both cases, extrusion-dependent G1 pruning of MCD-MCD interactions generally seems to entail melting of long-range inter-domain microcompartments into the broader canonical active compartments (Fig 5k). Notably, the differences in MCD-MCD interactions with respect to loop extrusion anchors observed 5 hours after mitotic release in RanGAP1-AID cells could be recapitulated in Nup93-AID cells confirming generality in DLD-1(Extended Data Fig. 5a-d). Taken together, we find that the extensive indiscriminate cCRE microcompartment formed during mitotic exit is pruned upon establishment of nucleocytoplasmic transport, particularly import of cohesin. We propose that cohesin-mediated loop extrusion, and the resulting TADs and CTCF-CTCF loops, impose constraints on which pair-wise MCD-MCD interactions remain, and which ones will melt into larger compartments (below).

### Interphase nuclei retain the capacity to form a cCRE microcompartment after establishment of the pruned G1 state

The cCRE microcompartment forms an indiscriminate grid of interactions genome-wide during telophase that is subsequently reduced to more specific and short-range contacts by early G1 (Fig. 5g). We sought to evaluate whether this pruned state could be reversed and the MCD contact grid re-established in the interphase nucleus. To this end, we performed Hi-C with cells depleted of RanGAP1 specifically in G1, after the normal interphase state had been acquired (Extended data Fig. 6a). Cells were first released from a prometaphase block and allowed to progress through mitosis in the presence of RanGAP1. Acute depletion of RanGAP1-AID was induced by addition of auxin 3.5 hours after prometaphase release, when cells are in G1. By 5 hours, the endpoint in previous experiments (Fig. 2), RanGAP1-AID was efficiently depleted as determined by Western Blotting (Extended data Fig. 6b) and cells were allowed to progress for an additional 5 hours into late G1. Hi-C analysis demonstrates that loss of RanGAP1 after early G1 enables the gain of long-range interactions between MCDs that are infrequent in control G1 cells (Extended data Fig. 6c) and indicates the capacity to re-form MCD contacts after pruning has occurred. The gain in MCD interactions coincides with reduced nuclear localization of cohesin upon G1-specific depletion of RanGAP1-AID (Extended Data Fig. 6d), suggesting that continuous nuclear transport capacity is required to maintain both normal cohesin levels in the nucleus and the pruned cCRE compartment.

**Fig. 6:**
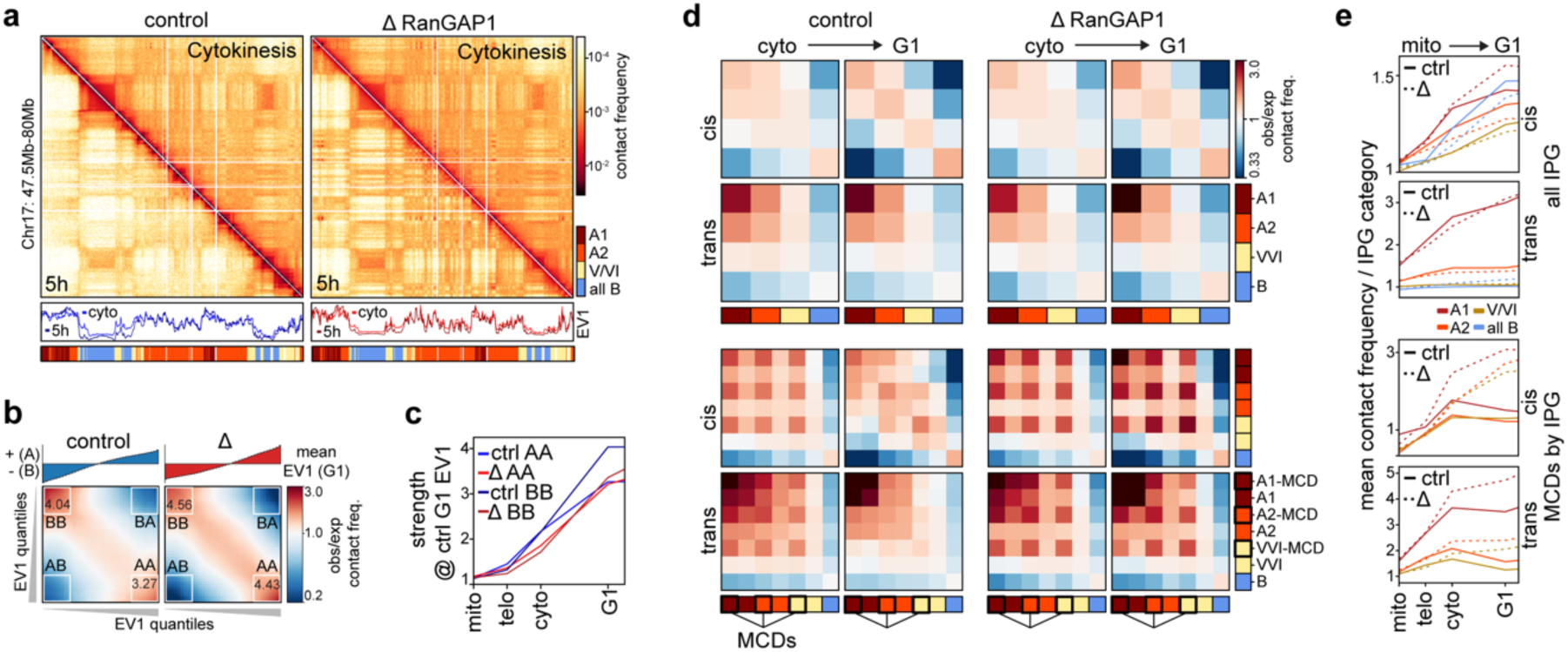
The post-mitotic microcompartment is resorbed by multiple distinct G1 compartments. a. Representative Hi-C interaction frequency maps at 50kb resolution (Chr7: 47.5-80 Mb) showing changes in genome organisation from cytokinesis (upper triangle) to G1 (t = 5h, lower triangle). Matched EV1 values and Interaction Profile Groups (IPGs) are shown. b. Saddle plots representing the segregation of active (A) and inactive (B) chromatin compartments in cis for control and RanGAP1-depleted cells 5h after mitotic release (early G1). The first eigenvector from each condition was used to rank 25 kb genomic bins into equal quantiles and the average interaction frequency between these ranked bins was normalized to the expected interactions to build the heatmap. Quantification of the average preferential A-A and B-B interactions for the top 20% strongest A and B loci are indicated. c. Quantification of average preferential A-A and B-B interaction strength in control and RanGAP1-AID-depleted cells demonstrating increased A/B segregation over the course of mitotic exit. Ranked bins designating A and B compartments were derived from the first eigenvector of the 5h control G1 cells at 25 kb resolution for comparison. d. Pairwise aggregate observed/expected contact frequency between DLD-1 IPGs showing enhanced homo-typic interactions in control and RanGAP1-AID-depleted cells from cytokinesis to early G1 (top). Aggregate observed/expected interactions between MCDs as a subset of each IPG demonstrate the cell cycle dynamics of a distinct micro-compartment in the presence or absence of RanGAP1 (bottom). e. Quantification of aggregate saddle plots in (d). Mean observed/ expected homo-typic interaction frequency is shown for all IPGs (top) or IPGs divided by the presence or absence of an MCD. Line colors are following the subcompartments shown in panel d.

Compared to control and prometaphase-depleted cells 10 hours after mitotic release, degradation of RanGAP1 during G1 results in a hybrid folding state. This mixed state is characterized by an increase in the size of cohesin-extruded loops, indicated by a shifted peak in *P*(*s*) curve (Extended Data Fig. 6e, vertical arrows), and the presence of both MCD-MCD interactions and convergent CTCF loops (Extended Data Fig. 6f).

Although loops stalled at pairs of CTCF sites appear unaffected, there is a quantitative reduction of extrusion flares emanating from CTCF motifs at convergent anchors (Extended Data Fig. 6g). These observations imply that continuous nuclear import is required for efficient cohesin-mediated loop extrusion in G1 but not the maintenance of existing positioned loops at pairs of CTCF sites. Loss of nuclear transport after G1 pruning has already occurred allows for an increase in average MCD interactions at all genomic distances compared to controls, both within and between looping domains as well as between chromosomes (Extended Data Fig. 6h). To a similar extent, interactions between active promoters and enhancers increase across length scales (Extended Data Fig. 6i). These data indicate a requirement for continuous active nuclear transport, and most likely loop extrusion, to maintain pruning of non-specific interactions between MCDs and suggest that these loci retain the capacity to form robust 3D contacts in interphase cells.

### The cCRE microcompartment is resorbed by multiple distinct active G1 subcompartments

We next investigated the relationship between the active cCRE microcompartment that appears transiently in telophase/cytokinesis and the global compartmentalization of chromosomes at the scale of compartments and subcompartments that persist in G1. We previously noted the use of Eigenvector decomposition to define A and B compartments from Hi-C data, where the first eigenvector (EV1) reflects the checkerboard patterns that are visually apparent in pairwise interaction maps. We determined EV1 for control and RanGAP1-AID depleted cells over the time course of mitotic exit and observed A/B compartmentalization in cytokinesis that becomes more pronounced in G1 (Fig. 6a). Compartment strength can be quantified by saddle plot analysis, where loci are ranked by EV1 such that homotypic A-A and B-B interaction frequencies can be compared to heterotypic A-B interactions ((Nora et al. 2017), Supplemental Methods, for example Fig. 6b). For both control and RanGAP1-AID depleted cells we observe very low compartmentalization scores in mitosis, and increasing strength as cells go through telophase, cytokinesis and G1 (Fig. 6c).

The absence of nuclear transport as cells progress to G1 affects the precise pattern of A/B compartmentalization along chromosomes (Supplementary Fig. 1c), evidenced by finer scale active domains captured in the first Eigenvector of the Hi-C matrix at 25 kb resolution (for example Fig. 6a, Extended Data Fig. 7a-b).

Furthermore, the segregation of these active and inactive chromatin compartments is enhanced when nuclear transport is not established during mitotic exit (Fig. 6b, Extended Data Fig. 7c). These observations are consistent with previous findings in Nipbl-depleted cells, where cohesin-dependent loop extrusion was effectively absent (Schwarzer et al. 2017). Despite these quantitative differences in A/B identity, transport competence during mitotic exit has minimal impact on the segregation, or strength, of compartments defined by control G1 Hi-C data (Fig. 6c, Extended Data Fig. 7d), supporting the intrinsic capacity for mitotic chromosomes to form epigenetically defined compartments, without a requirement for cytoplasmic factors.

The binary classification of active and inactive chromatin from Hi-C data is an oversimplification of genome compartmentalization (Imakaev et al. 2012) that can be expanded to include cell type-specific subcompartments of different types by various methods (Rao et al. 2014; Xiong and Ma 2019; Chen et al. 2018). One such approach clusters leading Eigenvectors of interchromosomal Hi-C contacts to reveal a set of subcompartments, or Interaction Profile Groups (IPGs), that correlate with distinct epigenetic signatures (Spracklin et al. 2023). To further investigate intrinsic compartmentalization, we employed IPGs that were previously defined in wild type DLD-1 cells (Scelfo et al. 2024) defining two distinct active (A1 and A2) and inactive (pooled to “all B”) chromatin subcompartments, as well as a transcriptionally intermediate group of genomic loci (V/VI). As expected for affinity driven compartmentalization, each of these four subcompartments show preferential homotypic interactions. These interactions increase as cells progress from cytokinesis to G1 in both control and RanGAP1-depleted cells, particularly within chromosomes (Fig. 6a,d-e (top)). We therefore conclude that compartmentalization at the finer scale of IPGs, as well as increased compartmentalization strength during G1 are entirely driven by determinants and factors associated with the telophase chromosome, with no requirement for factors from the G1 cytoplasm.

Finally, we asked whether the active cCRE microcompartment is equivalent to or contained within a specific subcompartment type. Surprisingly, MCDs are not enriched in any particular active subcompartment but are rather found with similar relative frequency at A1 (6.6%), A2 (6.7%), and V/VI (5.9%) and are largely absent from the pooled B compartment (0.3%). We show that interactions between MCDs are quantitatively and temporally distinct from homotypic interactions between corresponding subcompartments. Microcompartment domains were separated from their IPGs in order to compare the interactions of MCDs and non-MCD loci from each subcompartment (Fig. 6d-e (bottom)). In cytokinesis MCDs interact most frequently with other MCDs regardless of their subcompartment status. Importantly, the preference for MCD-MCD interactions is lost in control G1 cells when these domains instead interact with other loci, including non-MCDs, in accordance with their subcompartment status. Conversely, MCD-MCD interactions are already stronger in cytokinesis in the absence of RanGAP1, and continue to strengthen during G1, maintaining the preference for their microcompartmental interactions over contacts with other loci of the same subcompartment type. These results demonstrate that the cCRE microcompartment is distinct from subcompartments defined in control G1 cells. In control cells, MCDs initially interact regardless of subcompartment status but are resorbed in their corresponding subcompartment during mitotic exit. In nuclear import-deficient cells this resorption does not occur and the cCRE microcompartment continues to strengthen (Fig. 6e, Extended Data Fig. 7e).

### Identification of factors that require nuclear import for chromatin enrichment

Our data implicates nuclear import of cohesin in the loss of cCRE microcompartment contacts after cytokinesis. In order to better understand the chromatin state that gives rise to this unique compartmentalization, as well as cytoplasmic factors that normally modify this pattern, we set out to identify additional factors requiring nuclear import to access the genome at the end of mitosis. To this end, we employed stable isotope labeling by amino acids in cell culture (SILAC) (Ong et al. 2002) of import-deficient cells exiting mitosis for liquid chromatography-mass spectrometry (LC-MS) (Fig. 7a). Isotope-labeled cells were synchronized in prometaphase and either RanGAP1-AID or Nup93-AID was degraded 2 hours prior to mitotic release, as before. After entry to G1, 5 hours after release, nuclei were isolated by mechanical disruption (Supplemental Methods) and subjected to LC-MS.

**Fig. 7:**
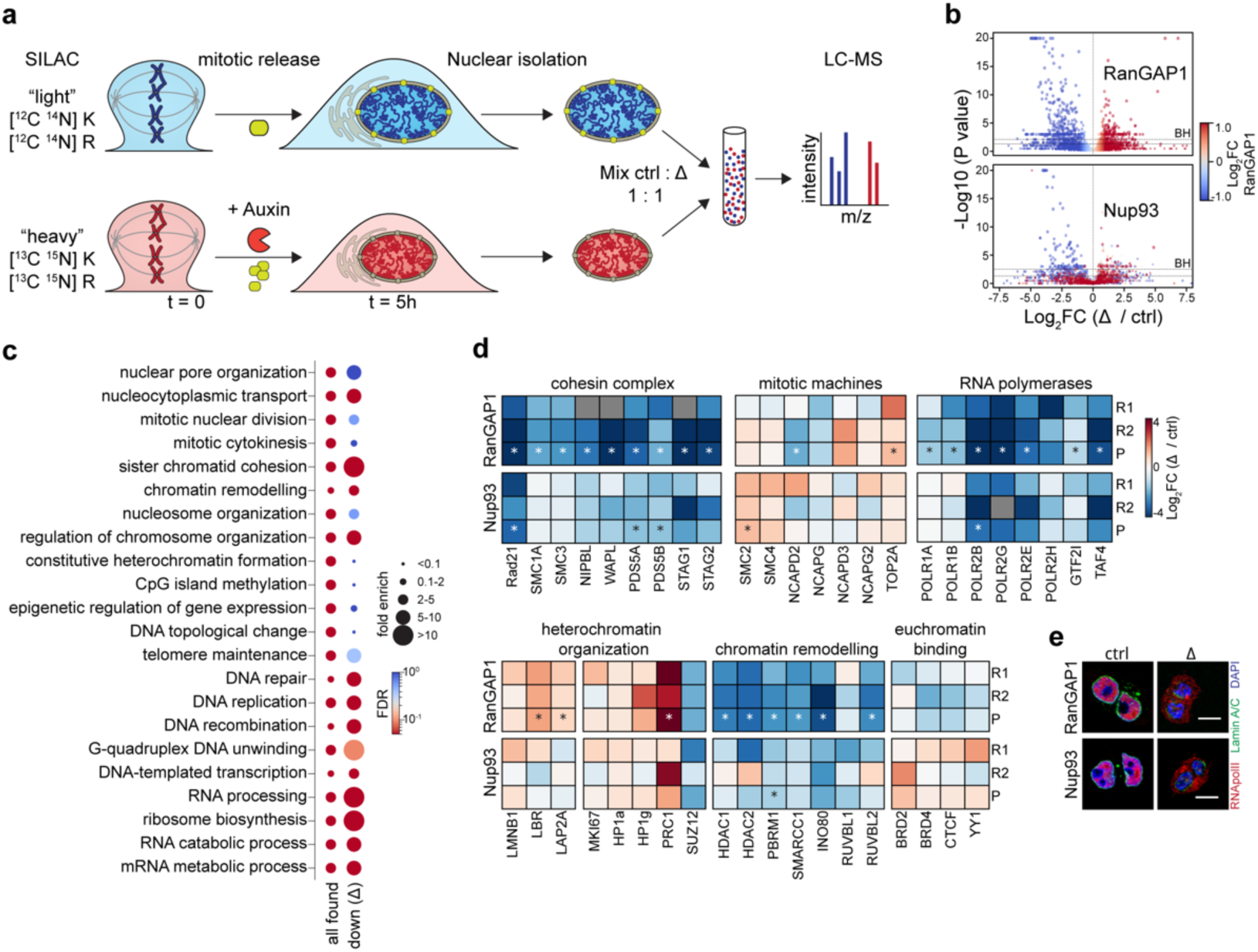
The nuclear proteome of transport deficient G1 cells. a. Experimental workflow for SILAC-based LC-MS quantification of nuclear proteome alterations found in cells entering G1 (t=5h) in the absence of either RanGAP1 or Nup93. Auxin-induced degradation is initiated 2h prior to nocodazole/mitotic release (t=0). b. Volcano plots for the consensus list of proteins identified in early G1 nuclear isolates from RanGAP1- and Nup93-AID cells. For each protein, the enrichment in auxin-treated vs. control cells (Log2-FC) is plotted against the BH-adjusted *p* value (p=0.05 and p=BH threshold are indicated). Two pooled replicates for heavy and light reversed experiments are shown and each protein is colored by the delta/ctrl ratio in the RanGAP1-AID. c. Gene ontology over-representation analysis for selected biological processes among all proteins identified in RanGAP1-AID and Nup93-AID control G1 nuclei (“all found”), or proteins reduced by at least 2-fold in the nuclei of both RanGAP-AID- and Nup93-AID-depleted cells (“down delta”). Dots are colored by the False Discovery Rate (FDR) and dot size indicates fold enrichment. d. Heatmaps representing the changes in the abundance of selected proteins involved in genome folding and function in RanGAP1-AID- or Nup93-AID-depleted G1 nuclei. The Log2-fold enrichment in auxin-treated vs. control nuclei was calculated separately for each individual replicate and for the pooled (reversed) LC-MS spectra. e. Representative immunofluorescence images of RanGAP1-AID or Nup93-AID control and depleted cells fixed 5h after mitotic release demonstrating the nucleo-cytoplasmic localisation of RNApolII (coloured red). Lamin A/C indicates the nuclear periphery. Scale bar represents 10 um.

As prominent MCD contacts are a strong feature of both RanGAP1 and Nup93 depletion, we focused on common changes to the nuclear proteome. Universal changes to protein composition in transport-deficient cells are almost exclusively under-represented factors that fail to localize to the nucleus as cells enter G1 (Fig. 7b, Supplemental tables 4-5). We classified these proteins according to annotated functions (Ashburner et al. 2000; Gene Ontology Consortium et al. 2023) and identified biological processes that are retained or lost in import-deficient nuclei (Fig. 7c). As expected, chromatin-associated processes are enriched in the proteomes of control isolated nuclei. Consistent with our observations of normal mitotic exit kinetics, the absence of nuclear transport does not consistently impact the presence of factors related to mitotic cell division, including proteins essential to mitotic chromosome condensation (Fig. 7d). We confirmed that the cohesin complex is completely absent in transport-incompetent nuclei. Other strongly depleted processes include DNA replication and repair, as well as RNA processing and ribosome biogenesis, the latter being consistent with the absence of nuclear speckles and nucleoli in depleted nuclei seen by immunofluorescence (Extended data Fig. 1e). Surprisingly, proteins involved in DNA-templated transcription were found to require nuclear import for access to the genome in early G1, confirmed by a dramatic reduction in components of the basal transcription machinery (Fig. 7d) and cytoplasmic localization of RNApolII by immunofluorescence (Fig. 7e) in both RanGAP1 and Nup93-depleted cells. Finally, although we found that changes to chromatin organization were relatively less consistent between depletions, we note that many chromatin remodeling enzymes are substantially absent from the nucleus in both RanGAP1 and Nup93 depletions, while structural components of heterochromatin and euchromatin may be less dependent on transport for nuclear localization in newly divided cells. We conclude that loop extrusion, transcription, and RNA processing likely do not occur, or occur to a dramatically reduced degree, in nuclei formed during mitotic exit in the absence of nuclear transport. Given that canonical chromatin compartments and the cCRE microcompartment form under these conditions, our data suggest that these active processes are mechanistically dispensable for compartmentalization during mitotic exit.

## Discussion

We propose the existence of two folding programs that specify interphase chromosome conformation as cells exit mitosis and enter G1. The first program is driven by factors that associate with chromosomes no later than telophase to drive genome-wide compartmentalization based on the epigenetic signatures of the previous cell cycle. This program includes and may even be driven by mitotically bookmarked cis-elements and trans-factors that form a prominent indiscriminate cCRE micro-compartment starting in telophase. Although inherited in the same way, the cCRE micro-compartment is distinct from conventional sub-compartments, both in composition and cell-cycle dependent dynamics. A second folding program starts after nuclear envelope formation and requires nucleocytoplasmic transport, implying that the relevant factors are inherited through mitosis in the cytoplasm. This second program includes cohesin, which drives the formation of interphase loops and TADs. It is intriguing that the two main known mechanisms of chromosome folding, compartmentalization and loop extrusion, are inherited through mitosis in two distinct and physically segregated ways.

The second (cytoplasmic) folding program modulates the first (chromosome-intrinsic) in several ways. First, the enrichment of cytoplasmic factors, most notably cohesin, generally reduces compartmentalization and leads to a loss of finer scale compartment interactions, consistent with previous observations in Nipbl-depleted cells (Schwarzer et al. 2017). Secondly, because most micro-compartment domains coincide with extrusion loop anchors (CTCF bound sites), the entry of cohesin into the nucleus likely modulates MCD-MCD micro-compartmental interactions directly.

The initially promiscuous cCRE micro-compartment appearing from telophase to cytokinesis is reduced to specific MCD interactions that reflect general rules of loop extrusion in early G1. Namely, long-range indiscriminate MCD interactions are depleted or pruned, and specific short-range interactions within TADs or at TAD boundaries are retained or even reinforced.

The interphase chromosome conformation is thus the result of the combined action of these two distinct folding programs. Our investigation into chromosome folding in nuclei deficient in nucleocytoplasmic transport from telophase onwards allowed us 1) to identify the existence of two distinct inherited folding programs; 2) to study the chromosome-intrinsic program in the absence of the cytoplasmic program; and 3) to demonstrate how their combined action determines the ultimate interphase folded state in normal unperturbed cells.

### A chromosome-intrinsic folding program

We describe a transient folding state during telophase and cytokinesis in which active cCREs display an intrinsic propensity to self-associate genome-wide, both in cis and in trans. We refer to this phenomenon as micro-compartmentalization in line with earlier work describing fine-scale compartmentalization of active elements (Goel et al. 2023). The extensive micro-compartment we observe during mitotic exit appears to be a transient folding state that is extensively modulated as cells enter and progress through G1, making it difficult to study. However, when nucleo-cytoplasmic transport is disrupted during mitotic exit (e.g., as in prometaphase depletion of RanGAP1 or Nup93), this micro-compartmentalized state not only persists but becomes increasingly strong as cells progress through the cell cycle. In this context, it was possible to characterize the loci involved in micro-compartmentalization in greater detail. We could identify strong constituent MCDs genome-wide explicitly from Hi-C data and discovered that they coincide with the active cCREs of the given cell type. We found that micro-compartmentalization is distinct from other types of sub-compartmentalization because MCDs are derived from all three active sub-compartments and display preferential homotypic interactions during cytokinesis that persist into G1 in the absence of nuclear transport. The fact MCDs are largely resorbed by subcompartments in control G1 cells could help to explain why they still interact with some frequency, as well as the capacity for chromosomes to restore microcompartment interactions when RanGAP1 is depleted in early G1. Although distinct, the cCRE micro-compartment and canonical A/B compartments and sub-compartments are all formed without import of factors from the cytoplasm, indicating that affinity-driven compartmentalization is generally mediated by factors that are stably associated with mitotic chromosomes, or rapidly recruited before telophase.

Several lines of evidence support the intrinsic nature of chromosome compartment formation. We found that the cCREs at strong MCDs are bookmarked during mitosis. Furthermore, histone modifications associated with active bookmarked loci (e.g., H3K4Me3, H3K72Ac) and more generally corresponding to subcompartments, are at least partly retained during mitosis. We note that there are conflicting reports regarding both the extent and precise cell cycle-dependent dynamics of histone modification deposition and conservation throughout mitosis (Behera et al. 2019; Zhiteneva et al. 2017; Kang et al. 2020). Nonetheless, active histone marks including H3K27Ac are thought to be fully present by telophase, consistent with a chromosome-intrinsic capacity for compartmentalization driven by epigenetic affinities. We found that the MCDs giving rise to stronger micro-compartments tend to overlap clusters of active gene-regulatory elements, suggesting that avidity contributes to interaction strength and implying that the affinities of the active elements themselves could drive the cCRE micro-compartment. It is possible that histone modifications are sufficient to mediate affinity-driven compartmentalization, although *in vitro* studies suggest that bridging factors, such as Brd4, are required for mediating interactions between loci marked with acetylated histones (Gibson et al. 2019). As proteins of the BET family, particularly Brd4, have been found to associate with mitotic chromosomes (Behera et al. 2019; Dey et al. 2009) and possess a capacity for phase separation (Sabari et al. 2018; Gibson et al. 2019), they represent one class of candidate bookmarked mediators of cCRE micro-compartmentalization. Previous work has shown that Brd2 and Brd4 contribute to active genome compartmentalization during interphase (Xie et al. 2022) and on condensin-depleted mitotic chromosomes (Zhao et al. 2024), respectively. However, as these proteins may function antagonistically, it remains to be determined whether bookmarked BET proteins mediate MCD-MCD interactions during mitotic exit.

Although compartmentalization is strongly correlated with gene expression patterns, we found that RNA polymerase II and most detectable components of the basal DNA-templated transcriptional machinery are excluded from the nucleus in RanGAP1-AID-depleted cells. The inheritance of transcriptional machinery through the cytoplasm, and its inability to access the genome in the absence of nuclear transport, confirms that chromatin compartmentalization driven by epigenetic states, and specifically bookmarked cCREs, is remembered from the previous cell cycle.

There is evidence to suggest that a burst of transcription occurs shortly after mitotic exit (Palozola et al. 2017; Hsiung et al. 2016) and we cannot exclude the possibility that early gene reactivation at bookmarked elements contributes to MCD interactions. However, our data suggests that ongoing transcription during G1 is dispensable for the formation of the cCRE micro-compartment. Similarly, compartmentalization has been linked to genome localization at speckles (Chen and Belmont 2019) and nucleoli (Bizhanova and Kaufman 2021) but the absence of these structures in nuclear transport-deficient cells did not adversely impact any level of global compartmentalization we examined.

Interactions between MCDs were not detected in prometaphase-arrested cells. Recent work has shown that acute degradation of condensin in prometaphase arrested cells leads to the formation of long-range chromosome interactions between loci marked by H3K27Ac (Zhao et al. 2024). As these cells are otherwise maintained in a mitotic state, it was proposed that the dense array of loops formed by condensins prevents an intrinsic capacity for mitotic chromosomes to form micro-compartments. In contrast to condensin-depleted prometaphase cells, we observe a loss of mitotic histone phosphorylation and changes to chromosome morphology and chromatin accessibility in RanGAP1-depleted G1 cells. However, as MCDs overwhelmingly overlap bookmarked open regions it is possible that the loss of condensins during telophase might then be sufficient to enable micro-compartment formation during mitotic exit.

### A folding program inherited through the cytoplasm

The post-mitotic nuclear landscape established in the absence of nucleocytoplasmic transport consists of partially decondensed chromosomes and lacks subnuclear organelles (e.g., speckles and nucleoli) as well as proteins involved in DNA metabolic processes (e.g. replication and repair) and RNA biosynthesis and processing (e.g., transcription and splicing). Most notably, cohesin is excluded from the nucleus and interphase extrusion-mediated chromatin structures, such as TADs and CTCF-CTCF loops, are absent. We find that a second folding program is inherited through the cytoplasm at the end of mitosis. The protein factors driving this program require nuclear import to access the genome and modulate the folded state generated by the first, chromosome-intrinsic, program.

The nuclear import requirement for cohesin accumulation at post-mitotic chromosomes, recently observed by quantitative and super-resolution microscopy (Brunner et al. 2024), segregates two modes of chromatin extrusion at the end of mitosis. This physical segregation is consistent with our previous work illustrating a chromosome folding intermediate during telophase, characterized by the absence of both condensin and cohesin loop extruding complexes (Abramo et al. 2019). The functional relevance of a loop-free chromosome conformation during mitotic exit remains unclear. It is possible that the time required to deacetylate cohesin so that it can re-associate with chromatin is simply longer than the time it takes to form a closed nuclear envelope. Alternatively, the simultaneous activity of multiple loop extruding complexes could be problematic, leading to complications when they collide or interfering with other processes that occur during anaphase and telophase including chromosome decondensation and removal of self-entanglements (Samejima et al. 2024; Hildebrand et al. 2024).

Our ATACseq analysis identifies a large set of putative cis-acting elements that become accessible during mitotic exit in a nuclear transport-dependent way. The cell-type specific G1 chromatin landscape thus relies not only on chromosome-intrinsic factors but also on those inherited from the cytoplasm. Bookmarked loci overlapping MCDs remain accessible across TF motifs throughout mitosis and eventually give rise to the cCRE microcompartment. However, it is also possible for bookmarked sites to be inaccessible during mitosis (e.g., CTCF in HeLa cells (Oomen et al. 2019)) and require active remodeling during G1. Accordingly, our proteomics analysis identified a number of chromatin remodeling complexes that rely on import for nuclear localization and may contribute to re-establishing the G1-specific architecture. Cohesin can be recruited anywhere along the genome, but loop extrusion patterns are sensitive to the presence and location of active cis-elements (Busslinger et al. 2017; Valton et al. 2022) and transcriptional machinery (Banigan et al. 2023). The second folding program inherited through the cytoplasm may therefore open distal regulatory sites and define a cell type-specific loop extrusion traffic pattern.

### Interplay between the two folding programs

The entry of cohesin, and other factors, into the nucleus modulates the chromosome-intrinsic micro-compartmentalization and sub-compartmentalization established during telophase and cytokinesis. Loop extrusion has a generally antagonistic effect on compartment segregation (Schwarzer et al. 2017; Rao et al. 2014) that is particularly obvious for the micro-compartmentalization of cCREs. Initial micro-compartmentalization is promiscuous: any pairwise interaction can occur, both in cis and in trans. As cytoplasmic factors enter the nucleus, long range (>1 Mb) MCD-MCD interactions are substantially reduced, or pruned. We have proposed that G1 pruning of indiscriminate MCD contacts is driven by cohesin-mediated loop extrusion, although roles for other factors cannot be excluded. The majority of MCDs contain CTCF sites that become anchors of extruded loops in G1. These MCDs retain short range interactions with other MCDs, possibly reinforced by loop extrusion. Pruning during mitotic exit ensures that MCD-MCD interactions start to obey rules of TAD formation such that interactions within extrusion domains are more likely to be retained than interactions between domains. Because the effect of pruning can be reversed by RanGAP1-depletion in G1, we conclude that interphase chromatin folding is the result of continuous interplay between intrinsic affinity-based compartmentalization and cytoplasmic inherited processes, such as loop extrusion.

### Functional relevance of the two programs

A chromosome-intrinsic folding program drives formation of a cCRE-anchored micro-compartment during the initial stages of mitotic exit. Indiscriminate MCD-MCD interactions appear to be affinity driven, occur in cis and in trans, and do not require loop extrusion. The formation of extrusion-independent promoter-enhancer contacts has been observed previously in different cell contexts and cell cycle stages (Goel et al. 2023; Lam et al. 2024; Calderon et al. 2022; Thiecke et al. 2020; Hsieh et al. 2022; Liu et al. 2021). During telophase and cytokinesis, we observe that the intrinsic capacity for previously active promoters and enhancers to interact gives rise to a comprehensive array of non-specific contacts. It is possible that this microcompartment serves to rapidly re-establish gene-regulatory contacts during mitotic exit. Acquisition of factors inherited from the cytoplasm, after the nuclear envelope has formed, restricts these interactions to short-range G1 loops that obey the rules of loop extrusion, a constraint that is consistent with a previous report of transient inter-TAD contacts during anaphase (Zhang et al. 2019). Furthermore, we find that microcompartment interactions giving rise to G1 loops are reinforced by loop extrusion, supporting the notion that de novo establishment and maintenance of gene-regulatory contacts during the cell cycle may operate by different mechanisms, in line with previous work (Lam et al. 2024). Cohesin-mediated loop extrusion, and its control by CTCF, prunes most longer-range micro-compartment interactions, which are reabsorbed into larger, more diffuse, sub-compartments. We propose that the chromosome-intrinsic folding program reveals a universal propensity for active gene-regulatory elements to interact through affinity-driven interactions. A second folding program superimposes a more deterministic logic through regulated cohesin-mediated loop extrusion that ensures the specific pairing of cCREs to ensure cell type-specific gene expression. Future experiments are required to understand the roles of individual factors as they contribute, through regulating cohesin or otherwise, to either of these folding programs and ultimately control chromosome conformation and gene regulation.

## Acknowledgements

We thank members of the Dekker lab, the Open2C community, especially Nezar Abdennur, for discussion on experiments and data analysis. We thank the UMass Electron Microscopy Core (Keith Reddig and Gregory Hendricks), the UMass Proteomics Core (Nadia Sultana and Scott Shaffer), the UMass FACS core (Tammy Krumpoch and Susanne Pechhold), the UMass Deep Sequencing Core (Ellie Kittler, Maria Zapp, Danielle Wilmot). We are grateful to Ross Kaufhold for help with constructing the MBP-mScarlet-NLS construct. This work was supported by grants from the National Human Genome Research Institute (HG003143 and HG011536 to JD). J.D. is an investigator of the Howard Hughes Medical Institute. V.A. and M.D. were supported by the Intramural Research Program of the *Eunice Kennedy Shriver* National Institute of Child Health and Human Development at the National Institutes of Health, USA (Intramural Project #Z01 HD008954). E.N. was supported by the Graduate Research Fellowship Program from the National Science Foundation.

## Conflicts of interest

Job Dekker is a member of the scientific advisory board of Arima Genomics, San Diego, CA, USA and Omega Therapeutic, Cambridge, MA, USA.

---

**Extended Data Fig.1:**
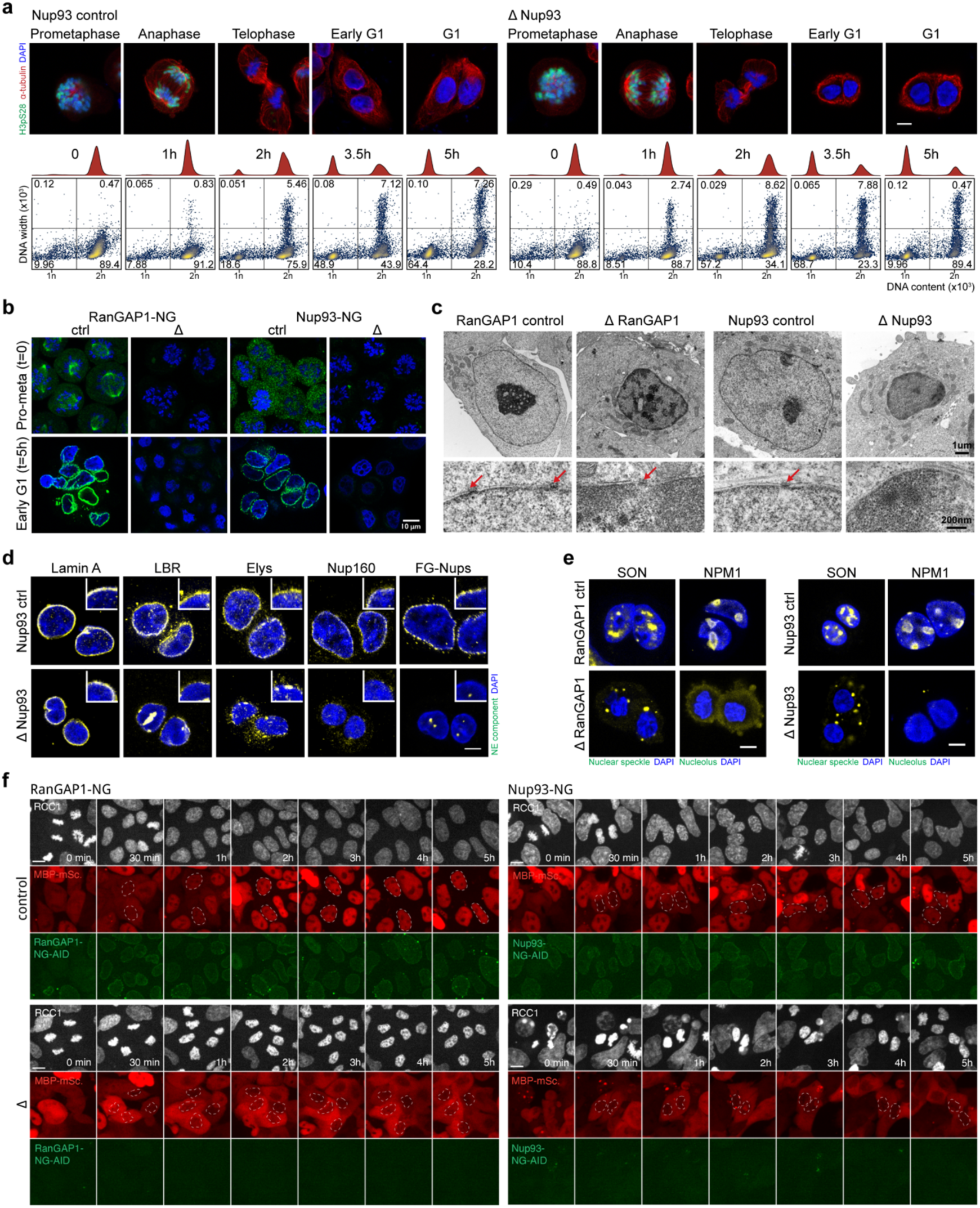
Acute depletion of RanGAP1-AID or Nup93-AID during mitotic exit enables the assembly of daughter nuclei isolated from the G1 cytoplasm. a. Representative immunofluorescence images for Nup93 control and depleted cells. Loss of Histone H3 serine 10 phosphorylation (H3pS10P) and DNA content at indicated time points indicates similar mitotic exit kinetics in the presence or absence of Nup93. Scale bar represents 5 um. Lower row of panels: DNA-content flow cytometry measurements for Nup93 control and depleted cells indicating similar mitotic exit kinetics in the presence or absence of Nup93. b. Fluorescent images of endogenous Neon Green-tagged RanGAP1-AID or Nup93-AID demonstrate efficient degradation of both proteins by 2 hours auxin treatment prior to mitotic release that is sustained into G1 in the presence of auxin. Scale bar represents 5 um. c. Transmission electron micrographs of RanGAP1-AID and Nup93-AID cells fixed at 5 hours reveal relatively small nuclei with hyper-condensed chromatin characteristic of post-mitotic depletion of either nuclear import factor. Arrows indicate nuclear pores embedded in the double lipid bilayers of the nuclear envelope and are not found in Nup93-depleted nuclei. d. Immunofluorescence images of Nup93 control and depleted cells 5 hours after mitotic release demonstrating presence of nuclear lamina (lamin A) and nuclear envelope (LBR) proteins as well as the DNA-binding nuclear pore complex protein, Elys, in the absence of Nup93. Structural (Nup160) and late associating (FG-Nups) nucleoporins are not found in Nup93-depleted nuclei. Scale bar represents 5 um and inset indicates a 5x magnification of the nuclear rim. e. Nuclear speckle (SON) and nucleolar (NPM1) resident proteins are mis-localized to the cytoplasm in RanGAP1-AID and Nup93-AID-depleted nuclei. Representative immunofluorescence images of cells fixed 5 hours after mitotic exit are shown. Scale bar represents 5 um. f. Representative time-lapse fluorescent images of RanGAP1-NG-AID or Nup93-NG-AID control and depleted cells blocked in G2 (Ro-3306) and released for 5h. Endogenously tagged RCC1-miRFP670 demarcating chromatin and the MBP-mScarlet-NLS import substrate are shown.

**Extended Data Fig. 2:**
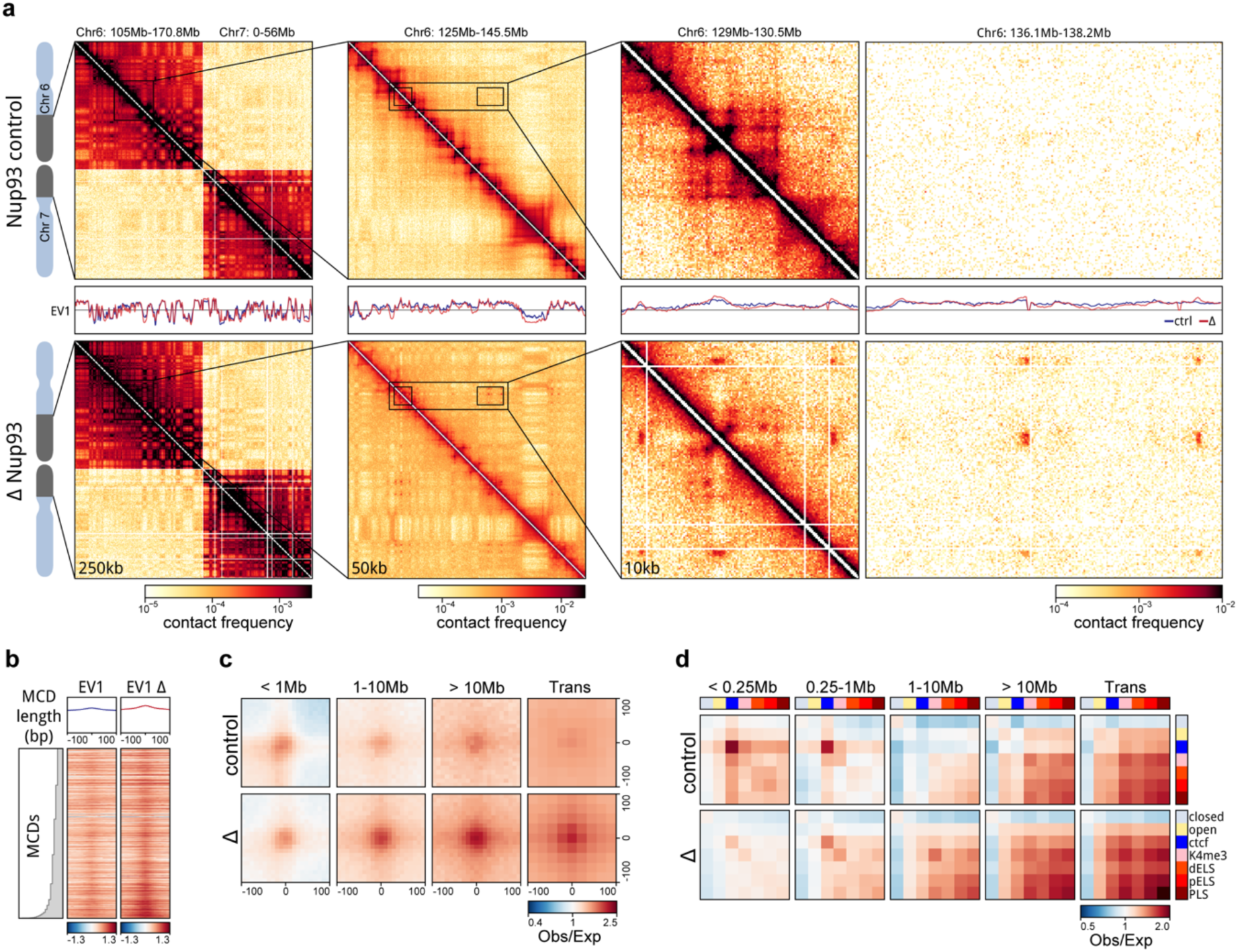
A chromosome-intrinsic capacity to form a kilobase scale cCRE compartment emerges upon depletion of Nup93 during mitotic exit. a. Representative Hi-C interaction frequency maps at 250kb (Chr 6: 105-170.8 Mb - Chr 7: 0-56 Mb), 50kb (Chr 6: 125 - 145.5 Mb), and 10kb (Chr 6: 129 -130.5 v. 136.1 - 138.2 Mb) resolution showing genome compartmentalisation in control and auxin-treated DLD-1 Nup93-AID cells released from prometaphase arrest for 5h. Matched first eigenvector (EV1) values for cis interactions are phased by gene density (A > 0). b. Heatmaps for all RanGAP1 MCDs centred on contact frequency summits and sorted by anchor length demonstrate differences in intra-chromosomal EV1 values derived from 10kb matrices in control and Nup93-depleted cells. b. Pairwise mean observed/expected contact frequency between all RanGAP1 MCDs projected in *cis* and *trans* showing enhanced interactions at all length scales and between chromosomes in Nup93-depleted G1 cells. c. Pairwise aggregate observed/expected contact frequency between cCREs assigned in control cells and subjected to hierarchical binning at 10kb resolution showing enhanced homo- and hetero-typic interactions between active promoters and enhancers in Nup93-depleted cells compared to controls at multiple genomic distances in *cis* and in *trans*.

**Extended Data Fig. 3:**
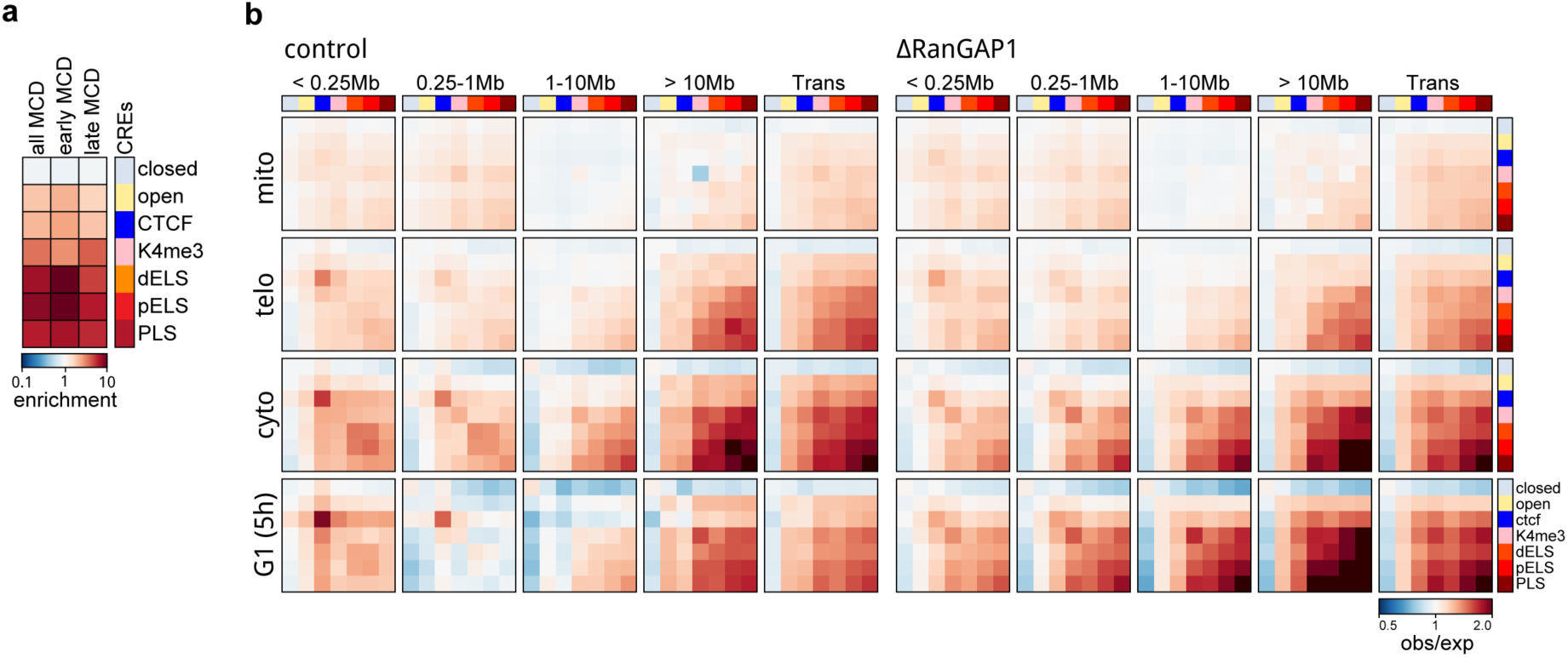
Cis-regulatory elements form a transient microcompartment during mitotic exit. a. Relative fold enrichment of control cCREs at early (cytokinesis) or late (G1-specific) MCD anchors demonstrating the predominance of active promoters and enhancers at microcompartment domains. b. Pairwise aggregate observed/expected contact frequency between cCREs assigned in control cells and subjected to hierarchical binning at 10kb resolution showing transient enhanced homo- and hetero-typic interactions between active promoters and enhancers at multiple genomic distances in *cis* and in *trans* during mitotic exit.

**Extended Data Fig. 4:**
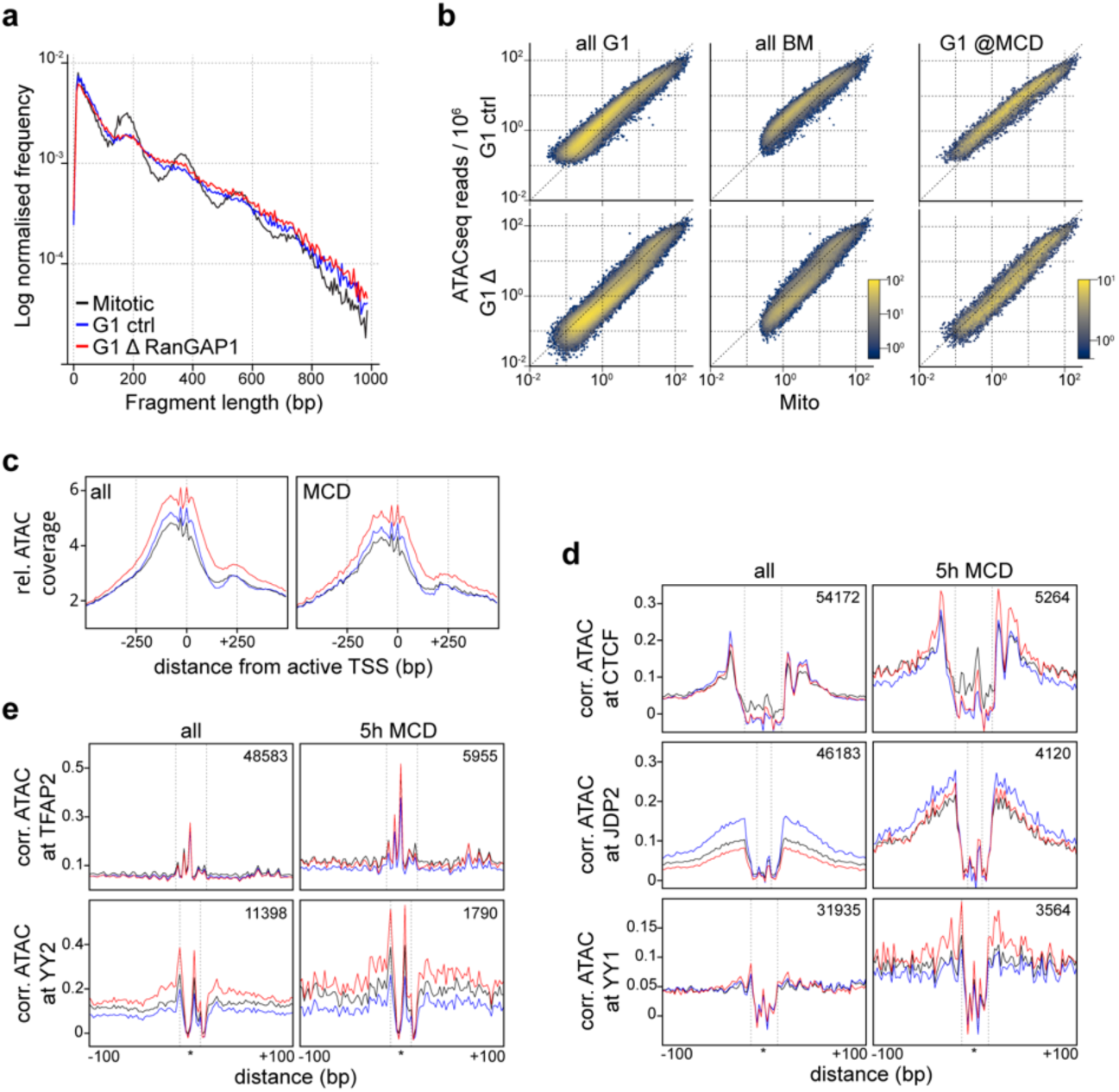
Chromatin accessibility at microcompartment domains during mitosis and G1. a. Relative fragment length distributions of ATACseq reads indicating regular nucleosome positioning in mitotically arrested cells and more dynamic G1 architecture in both control and RanGAP1-depleted cells released for 5h. b. Comparison of G1 and prometaphase ATACseq read coverage at all control G1, bookmarked, or MCD-overlapped G1 peaks demonstrating a common shift towards higher coverage peaks in all conditions at the MCDs. c. Aggregation plots of prometaphase, G1 control, or RanGAP1-depleted G1 ATACseq signal centred at active promoters (TSS) (FPKM >1) displaying genome-wide (left) and MCD-overlapped (right) averages. TSS are oriented in the same forward direction and ATACseq signal is binned to 5bp resolution and normalised to the mean frequency of the 1 kb upstream window. d. Aggregation plots of Tn5 bias-corrected (Tobias) prometaphase, G1 control, or RanGAP1-depleted ATACseq signal centred at bookmarked (TFAP2) or G1 control-specific (JDP2) example TF motifs (d) or motifs with high RanGAP1 depletion footprints (CTCF, YY1 and YY2) (e) indicating relative differences in footprint protection at genome-wide G1 or MCD-overlapped peaks. Motifs are oriented in the same forward direction and ATACseq signal is binned to 2bp resolution.

**Extended Data Fig. 5:**
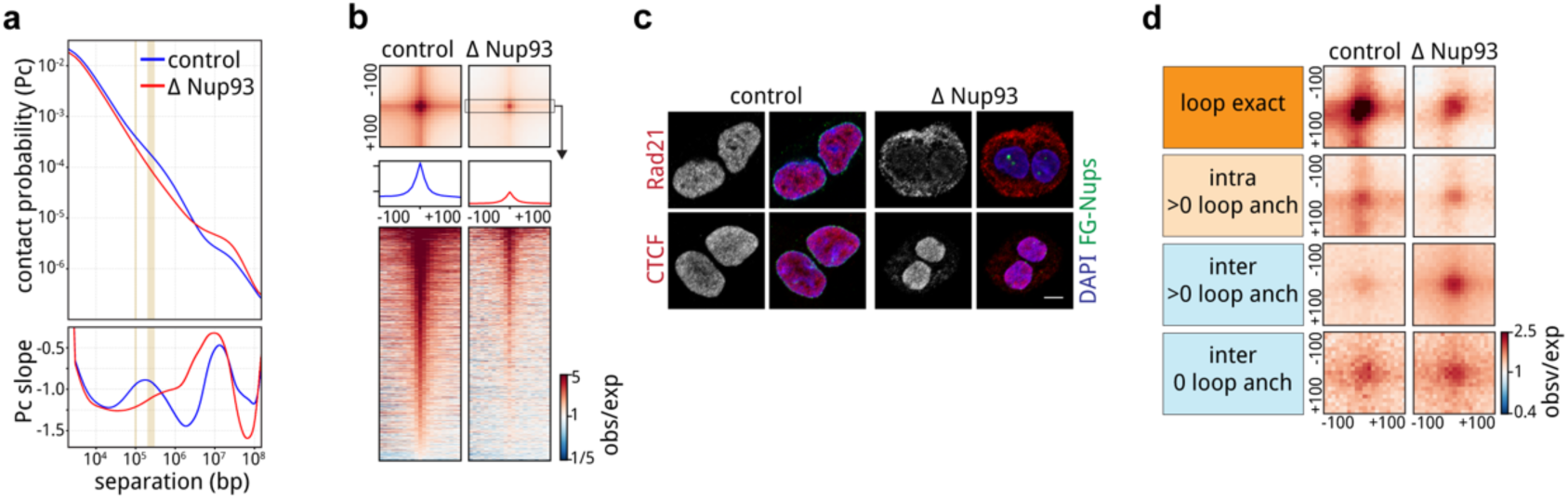
Microcompartment pruning generally requires nuclear transport in G1. a. P(s) and derivative P(s) plots for Hi-C data from FACS-sorted early G1 (t = 5h) control and Nup93-depleted cells. b. Pairwise mean observed/expected Hi-C contact frequency between convergent CTCF loops identified in pooled interphase RanGAP1-AID control Hi-C data demonstrating Nup93-dependent looping interactions in early G1 (t = 5h). Average signal for three central 10kb bins across the 200kb CTCF motif-centred window and stack-ups sorted by G1 loop strength are shown. c. Representative immunofluorescence images of Nup93-AID control and depleted cells fixed 5h after mitotic release demonstrating the nucleo-cytoplasmic localisation of the cohesin complex subunit, Rad21, and boundary transcription factor, CTCF. Scale bar represents 5 um. d. Pairwise mean observed/expected contact frequency between MCDs in early G1 (t = 5h) in control and RanGAP1-depleted cells. MCD-MCD contacts are categorised by the presence of a convergent CTCF loop or loop anchor (> 0) and the looping domain status of the constituent MCDs, as indicated.

**Extended Data Fig 6:**
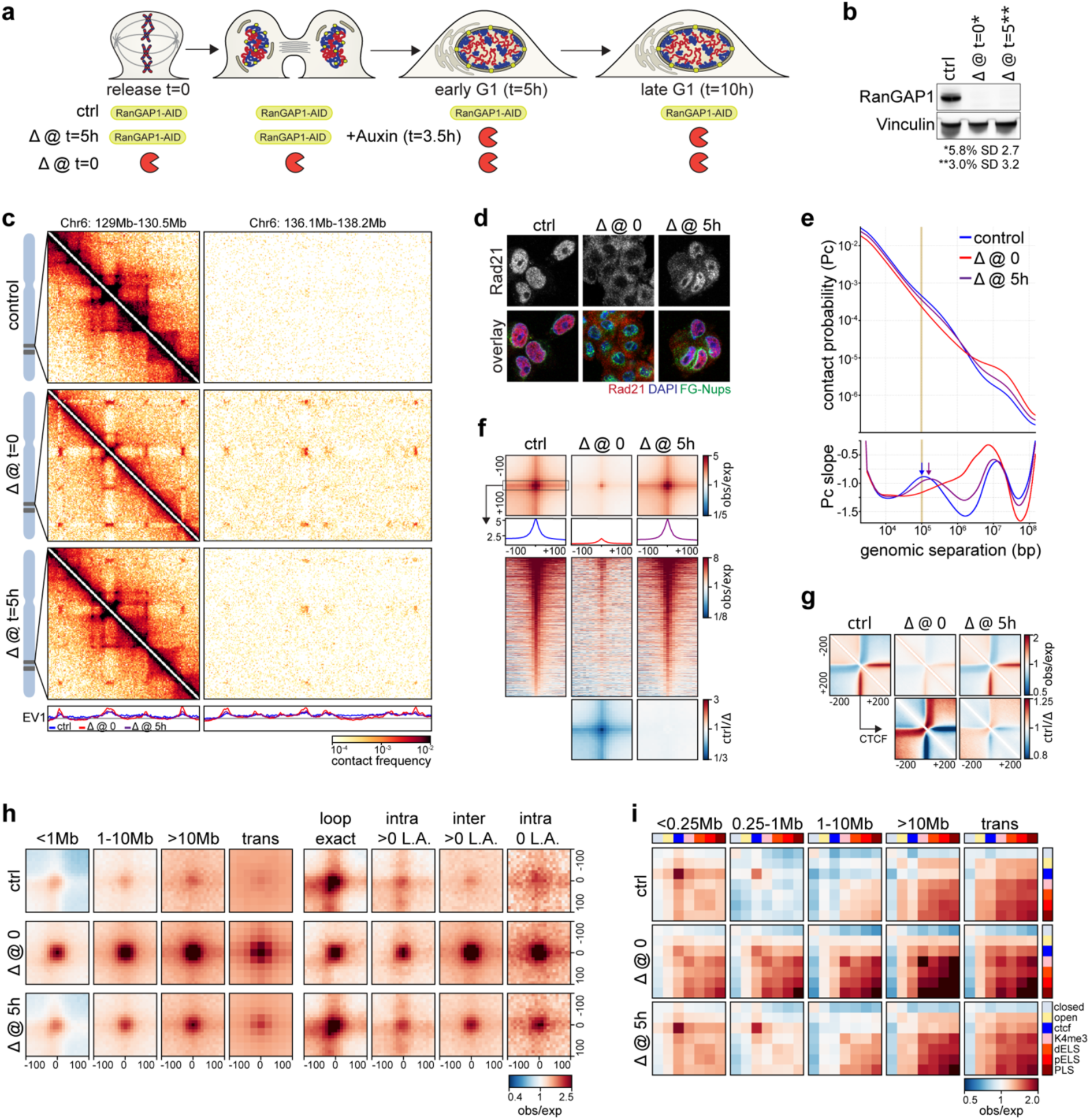
The capacity for genome-scale microcompartment formation is retained in G1 cells. Experimental workflow for the depletion of RanGAP1-AID in early G1. Auxin-induced degradation is initiated 3.5h following nocodazole release to ensure the absence of RanGAP1 protein by t = 5h. Control, Mitotic degraded (t = 0), and early G1-degraded (t = 5h) RanGAP1-AID cells are fixed for Hi-C 10h after prometaphase release in late G1. a. Representative western blot images of whole cell lysates derived from control and mitotic (t = 0) or G1 (t = 5h) auxin-treated DLD-1 cells 10h after mitotic release showing efficient depletion of RanGAP1. Mean protein levels normalised to vinculin for depleted lysates relative to untreated controls are shown for 3 replicates with standard deviation (SD). b. Hi-C interaction frequency maps at 10kb (Chr 6: 129 - 130.5 & 136.1 - 138.2 Mb) resolution showing genome compartmentalisation in RanGAP1-AID control and mitotic (t = 0) or G1 (t = 5h) depleted cells released from prometaphase arrest for 10h. Matched first eigenvector (EV1) values for cis interactions are phased by gene density (A > 0). c. Representative immunofluorescence images of RanGAP1-AID control and mitotic (t = 0) or early G1-depleted (t = 5h) cells fixed 10h after mitotic release demonstrating the nucleo-cytoplasmic localisation of the cohesin complex subunit, Rad21. Scale bar represents 5 um. d. P(s) and derivative P(s) plots for Hi-C data from synchronised RanGAP1-AID control and mitotic (t = 0) or early G1-depleted (t = 5h) cells fixed 10h after mitotic release. Vertical goldenrod line indicate the average loop size in control cells. Arrows highlight that in RanGAP1-AID-depleted cells the average loop size increases. e. Pairwise mean observed/expected Hi-C contact frequency in late G1 (t = 10h) between convergent CTCF loops, demonstrating retained looping interactions when RanGAP1-AID is degraded in early G1 (t = 5h). Average signal for three central 10kb bins across the 200kb CTCF motif-centred window and stack-ups sorted by G1 loop strength are shown. The ratio of observed/expected interaction frequency for depleted vs control cells is shown. f. Observed/expected Hi-C interaction pile-ups centred on forward-oriented CTCF motifs that overlap loop anchors are plotted in a 400kb window at 10kb resolution. The ratio of observed/expected interaction frequency for depleted vs control cells is shown. g. Pairwise mean observed/expected contact frequency between MCDs 10h after prometaphase release, in control and mitotic (t = 0) or early G1-depleted (t = 5h) RanGAP1-AID cells. MCD-MCD contacts are categorised by distance or the presence of a convergent CTCF loop or loop anchor (“> 0”) and the looping domain status of the constituent MCDs, as indicated. h. Pairwise aggregate observed/expected contact frequency between cCREs assigned in control cells and subjected to hierarchical binning at 10kb resolution showing enhanced homo- and hetero-typic interactions between active promoters and enhancers at multiple genomic distances in *cis* and in *trans* in late G1 (t = 10h) for G1-depleted (t = 5h) RanGAP1-AID cells.

**Extended Data Fig. 7:**
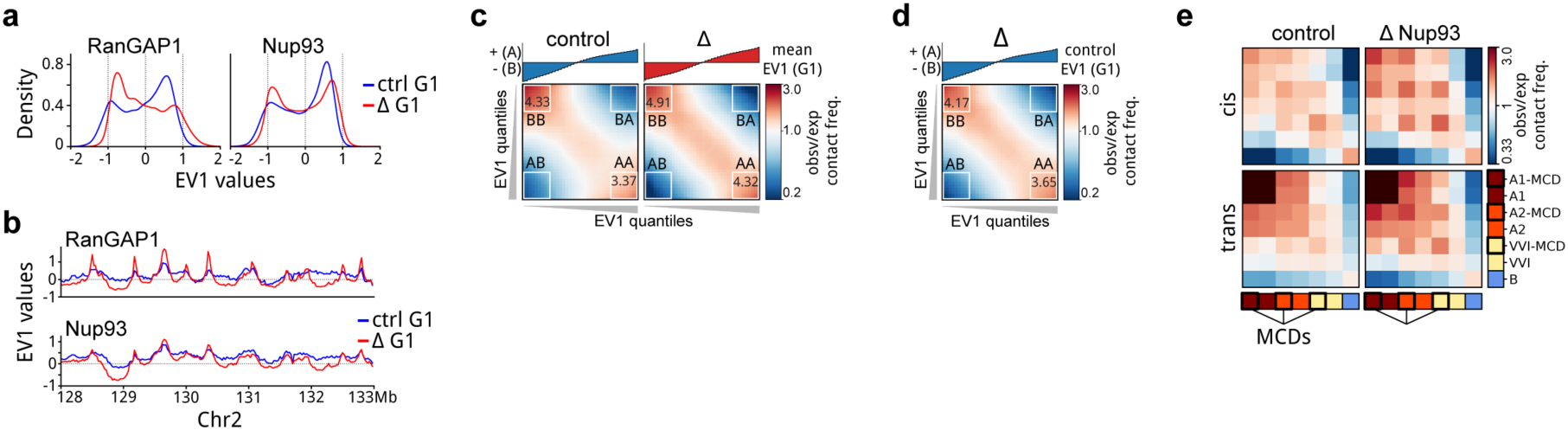
A distinct microcompartment is formed in the absence of RanGAP1 or Nup93. a. Distributions of EV1 values from Eigenvector decomposition of 25kb binned Hi-C data from control and auxin-treated RanGAP1-AID and Nup93-AID cells in G1. b. Representative examples of 25kb EV1 tracks from the Hi-C data of control and RanGAP1-AID- or Nup93-AID-depleted cells in early G1. c. Saddle plots representing the segregation of active (A) and inactive (B) chromatin compartments in cis for control and Nup93-AID-depleted cells 5h after mitotic release (G1). The first eigenvector from each condition was used to rank 25 kb genomic bins and quantification of the average preferential A-A and B-B interactions for the top 20% strongest A and B loci are indicated. d. Saddle plots representing the segregation of active (A) and inactive (B) chromatin compartments defined in control cells at 25kb resolution and plotted for Nup93-AID-depleted cells 5h after mitotic release (G1). Quantification of the average preferential A-A and B-B interactions for the top 20% strongest A and B loci are indicated. e. Pairwise aggregate observed/expected contact frequency between IPGs derived from DLD-1 cells and further categorized by the presence or absence of MCDs showing enhanced homo-typic interactions between MCDs in Nup93-AID-depleted cells in early G1.

