## Supplemental Methods for "Interphase chromosome conformation is specified by distinct folding programs inherited via mitotic chromosomes or through the cytoplasm"

**Supplementary methods**

**Genome modifications**

The CRISPR/Cas9 system was used to endogenously target the RanGAP1, RCC1 (Aksenova et al. 2022, 2020), NUP93 (Regmi et al. 2020) and AAVS1 (Chu et al. 2015) genes. With the exception of nuclear import assays, all experiments described here employed human colorectal adenocarcinoma DLD-1 cells (ATCC CCL-221) expressing either RanGAP1 or Nup93 homozygously tagged with NeonGreen and an Auxin-Inducible Degron, Infra-Red protein (IFP)-tagged RCC1 and Tir1, as described previously. We refer to these cell lines as RanGAP1-AID and Nup93-AID.

For cell lines used in nuclear import assays, the sequences of MBP and mScarlet, were amplified by PCR from pMAL (NEB) and pmScarlet_alphaTubulin_C1 (Addgene, #85045), respectively. The NLS sequence was synthesized, and all fragments were inserted by Gibson reaction (E2611S, NEB) into the MCS of AAVS1_Puro_PGK1 vector (Addgene, #68375) through replacement of 3xFlagTwinStep-Tag. MBP-mScarlet-NLS was inserted into the AAVS1 locus in RanGAP1-AID and NUP93-AID DLD-1 cell lines.

**Cell culture and cell cycle synchronization**

Modified DLD-1 cells lines (RanGAP1-AID, Nup93-AID, RanGAP1-AID/MBP-NLS, and Nup93/MBP-NLS) were cultured at 37 °C with 5% CO2 in in DMEM, high glucose, GlutaMAX supplement with pyruvate (Gibco, 10569010) with 10% FBS (Gibco, 16000069) and 1% penicillin-streptomycin (Gibco, 15140122). Cells were synchronized in prometaphase of mitosis using a standard double thymidine-nocodazole block protocol. All incubations and washes were carried out in pre-warmed solutions at 37 °C. Cells were seeded at low density in DMEM containing 4mM Thymidine (Sigma, T1895) and incubated for 17 hours, released (after washing 2x with PBS) for 7 hours in fresh medium to replicate the genome and enter mitosis, and incubated with 4mM Thymidine for an additional 17 hours to obtain synchronous populations blocked in S-phase. After the second thymidine block, cells were released (after washing 2x with PBS) in fresh medium for 5 hours followed by an additional 5h incubation in the presence of 50ng/ml Nocodazole (Sigma, M1404) to accumulate in prometaphase. Degradation of RanGAP1- and Nup93-AID- was achieved in prometaphase-arrested cells by addition of 1mM 3-Indoleacetic acid (IAA, auxin) (Sigma Aldrich, 15148). After 2 additional hours of incubation, control and depleted cells were either collected for subsequent analyses described below or released into fresh medium to progress through mitosis for 1.25h (telophase), 1.5h (cytokinesis), 5h (early G1), or 10h (late G1). For the G1 depletion of RanGAP1, auxin was added 3.5h after Nocodazole release to ensure degradation by 5h, followed by incubation for an additional 5h into late G1.

**Import assay of MBP-mScarlet-NLS in AID-tagged cell lines**

DLD-1 RanGAP1-AID and NUP93-AID cell lines, expressing MBP-mScarlet-NLS, were grown on 4-well glass bottom chambers (Ibidi) and synchronized with CDK1 inhibitor (Ro-3306, Selleckchem) for 16 h. After 15 h in the CDK1 inhibitor, cells were treated with fresh media or media containing 1 mM auxin for 1 h in the presence of the CDK1 inhibitor. Cells were washed three times with fresh media and released in regular media or media containing 1 mM auxin.

Cells were imaged on the Eclipse Ti2 inverted microscope (Nikon), equipped with a spinning disk confocal system (UltraVIEW Vox Rapid Confocal Imager; PerkinElmer) and controlled by Volocity software (PerkinElmer) utilizing Nikon PlanFluor 40x/1.3 oil immersion objective lens. Cells were imaged in FluoroBrite DMEM (ThermoFisher) media. The microscope was equipped with a temperature-, CO_2_- and humidity-controlled chamber that maintained a 5% CO_2_ atmosphere and 37°C.

RanGAP1-NG, NUP93-NG, MBP-mScarlet-NLS, and RCC1iRFP670 fluorescent protein signals were excited with a 488-nm (20-24% of power, 200-250 ms exposure), 568-nm (5% of power, 50-60 ms), 640-nm (100% of power, 150-200ms exposure) laser lines and binning set to 2x, respectively. A series of 6 z-slices 2 µm optical sections were acquired every 2 min and monitored for 6 hours for 3 independent fields. Images were captured and analyzed using Volocity (PerkinElmer) and Image J (National Institutes of Health) software, respectively. Images represent a single z-stack. Scale bar 10 um.

**Nuclear import image analysis**

The intensity of the MBP-mScarlet-NLS in three independent ROIs (5 x 5 pixels) was measured every 10 min in the nucleus and the cytoplasm. To calculate relative nuclear intensity, we used the following equation:

*Rel Int.Nu = Avg Int.Nu - Avg Int.B / ( Avg Int.Nu - Avg Int.B ) + ( Avg Int.Cyt - Avg Int.B)*,

where *Avg Int.Nu* is the average intensity of MBP-mScarlet-NLS in the nucleus, *Avg Int.Cyt* is the average intensity of MBP-mScarlet-NLS in the cytoplasm*,* and *Avg Int.B* is the average background intensity of MBP-mScarlet-NLS in cell-free area.

Relative nuclear to cytoplasmic intensity of MBL-mScarlet-NLS was plotted. Data are expressed as individual values analyzed every 10 min for eighth control or auxin-treated cells of two independent fields of one experiment. Assessment of MBL-mScarlet-NLS import has been performed in two independent experiments.

**Western blotting**

Adherent cells were dissociated with accutase and equal numbers of cells were lysed with 2x Laemmli Buffer (60mM Tris pH6.8, 10% glycerol, 2% SDS, 100mM DTT) by boiling for 10 minutes. Proteins were separated on 4-12% NuPage Bis-Tris gels using 1x MES running buffer for 45 minutes at 175V in an Invitrogen XCell Sure-Lock mini gel and blotting system. Gels were transferred to 0.2um nitrocellulose membrane in Pierce 10X Western Blot Transfer Buffer, Methanol-free (Thermo Fisher Scientific, 35040), by running for 1 hour at 30V. For immunoblotting, membranes were blocked with 5% milk in TBS-T (1x TBS + 0.1% Tween-20) for at least 1 hour at room temperature. Primary antibodies were diluted in block buffer and incubated overnight at 4°C and HRP-linked secondary antibodies for 1h at room temperature. Blots were developed and imaged using SuperSignal West Dura Extended Duration Substrate (Thermo, 34076) and a Bio-Rad ChemiDoc.

Primary antibodies: mouse anti-RanGAP1 (OTI1B4, Novus Biologicals, NBP2-02623), mouse anti-Nup93 (F-2, Santa Cruz Biotechnology, sc-374400), and rabbit anti-vinculin (EP18185, Abcam, ab129002). Secondary antibodies: goat anti-mouse IgG-HRP (Cell Signaling 7076), goat anti-rabbit IgG-HRP (Cell Signaling 7074).

**Immunofluorescence**

Immunofluorescence was performed using standard methods. For early G1 IF, cells were released from prometaphase onto glass coverslips and fixed with 4% paraformaldehyde in PBS for 30 minutes at room temperature. For mitotic release experiments, cells were first fixed and then concentrated onto glass coverslips using an Epredia Cytospin at 800 rpm. Fixed samples were incubated with PBS containing 2% Triton-X-100 for 1h prior to primary and secondary antibody staining in PBS containing 0.1% TX100, for 3h or 1h, respectively. After staining with 10 ug/ml DAPI (4′,6-diamidino-2-phenylindole) in PBS for 10 minutes at room temperature, coverslips were mounted in Vectashield antifade medium (Vector Labs, H-1000-10) for confocal imaging. Primary antibodies: mouse anti-alpha-tubulin (Sigma T6199), rabbit anti-histone H3pS28 (Abcam, ab5169), rabbit anti-Lamin B-receptor (Abcam, ab32535), rabbit anti-Lamin A (Abcam, ab26300), mouse anti-Elys (BioMatrix research, BMR00513), rabbit anti-Nup160 (Abcam, ab73293), mouse anti-Mab414 (Abcam, ab24609), rabbit anti-SON (Thermofisher, PA5-65107), mouse anti-NPM1 (Thermofisher, 60096-1), rabbit anti-Rad21 (Abcam, ab154769), rabbit anti-CTCF (Cell signaling, 2899), rabbit anti-RNAPolII pS2 (Abcam, ab5095). Secondary antibodies: goat anti-mouse IgG H+L Alexa Fluor 488 (Abcam, ab150113), goat anti-mouse IgG H+L Alexa Fluor 568 (Abcam ab175473), goat anti-rabbit IgG H&L Alexa Fluor 488 (Abcam ab15007), goat anti-rabbit IgG H&L Alexa Fluor 568 (Abcam, ab175471).

**Confocal microscopy**

Images were acquired using a Nikon A1 point-scanning confocal microscope with GaAsP detectors (488 and 561 lasers) or a high sensitivity MultiAlkali PMT (405 laser) and Apo TIRF, N.A. 1.49, 60x oil immersion objective (Nikon). For chromatin volume estimation, fixed DAPI-stained cells were imaged in 40-60 consecutive 0.2 um z-slices. The 3D volume of DAPI-containing signal was used as a proxy for chromatin volume and measured using the “3D Objects Counter” Plugin (Bolte and Cordelières 2006) as part of Fiji software (Schindelin et al. 2012) on pre-filtered (Gaussian, sigma = 2) image stacks.

**Flow cytometry**

Cells were collected at various points of mitotic exit. Adherent cells were dissociated with accutase (ThermoFisher Scientific, A11105-01) and pooled with non-adherent collected cells in order to assess the entire population. To assess the cell-cycle profile (DNA content), cell pellets were resuspended in 200ul PBS and fixed with 800 µl of cold 100% ethanol. Cells were stored at −20 °C for at least 24 h. Approximately 1 million fixed cells were stained with 50 ug/ml propidium iodide (PI) (Thermo, P1304MP), diluted in 1 ml PBS containing 50 ug/ml RNaseA (Roche, 10109169001) and 0.1% Saponin, for 1 hour at room temperature. After staining, cells were spun and pellets were resuspended in 1 ml of PBS and passed through a 35 *u*m filter (Falcon 352235). Flow cytometry was performed on a MACSQUANT set-up. Analysis was performed using FlowJo software (v10) and plots reflect populations gated for debris but not doublets.

**Hi-C fixation and fluorescence-activated cell sorting (FACS)**

Cells were collected at various points of mitotic exit and fixed for Hi-C 3.0 (Lafontaine et al. 2021) with a few modifications to facilitate cell sorting. Adherent cells collected 5 and 10 hours after prometaphase release, were dissociated with accutase. Prometaphase (t=0) and early (t=1.25-1.5h) released cells were directly collected by shake-off. Cell suspensions were pelleted and treated with accutase for an additional 5 minutes at room temperature to prevent aggregation and washed with HBSS (Thermofisher, 14025134). Fixation proceeded first with 1% formaldehyde (Fisher, BP531-25) in HBSS for 10 minutes, which was quenched with 0.125M Glycine for 5 minutes at room temperature and 15 minutes on ice. Next, cells were fixed with 3mM disuccinimidyl glutarate (DSG) in PBS for 40 minutes rotating at room temperature, followed by a second quenching with 0.125M Glycine. Fixed cells were washed twice with PBS + 0.1% BSA and snap-frozen in liquid nitrogen prior to staining for Fluorescence-activated cell sorting (FACS).

In order to sort cells by DNA content approximately 10 million fixed cells were stained with 50 ug/ml PI, diluted in 5 ml PBS containing 50 ug/ml RNaseA and 0.1% Saponin, for 1 hour at room temperature. Cells were then spun and washed with PBS prior to resuspension in 2 ml PBS + 0.1% BSA and passage through a 35 *u*m filter. Propidium iodide-stained cell suspensions were sorted on a BD FACS Melody using the 561 nm laser for FSC, SSC, and PI. All populations were gated based on FSC/SSC to eliminate cell debris and cells sorted for either prometaphase (4n) or G1 (2n) DNA content were also subject to doublet discrimination. To enrich for telophase or cytokinesis, cells fixed 1.25 and 1.5 hours after mitotic release, respectively, were sorted based on doubled PI signal (DNA content) area and width. All sorted cells were collected in PBS containing 1% BSA and washed twice in PBS prior to snap-freezing.

**Hi-C**

Chromosome conformation capture was performed as previously described (Lafontaine et al. 2021) with some modifications. Synchronized cells were crosslinked with 1% formaldehyde and 3mM DSG and enriched in specific cell cycle stages by FACS, as described above. 1-5 million sorted cells were collected and snap-frozen for storage at -80 °C prior to lysis. After lysing cells and digesting chromatin with 400U each of DpnII and DdeI (NEB, R0543M and NEB, R0175) overnight, DNA ends were labeled with biotinylated dATP (LifeTech, 19524016)) using 50 units Klenow DNA polymerase (NEB, M0210). Blunt-end ligation was performed with 50 units T4 Ligase (Life Technologies, 15224090) at 16 °C for 4 h, followed by reverse crosslinking with 400 μg/ml proteinase K (ThermoFisher, 25530031) at 65 °C overnight. DNA was purified using phenol/chloroform extraction and ethanol precipitation, and concentrated on a 30 kDa Amicon Ultra column (EMD Millipore, UFC5030BK). Biotin was removed from unligated ends in 50 μl reactions using 50 units of T4 DNA polymerase (NEB, M0203) per 5 mg of DNA. Following DNA sonication (Covaris S220) and SPRI bead size fractionation to generate DNA fragments of 100–300 bp, DNA ends were repaired using 7.5 units of T4 DNA polymerase, 25 units of T4 polynucleotide kinase (NEB, M0201) and 2.5 units of Klenow DNA polymerase (NEB, M0210). Libraries were enriched for ligation products by biotin pulldown with MyOne streptavidin C1 beads (Invitrogen, 65001). To prepare for sequencing, A-tailing was performed using 15 units of Klenow DNA polymerase (3′–5′ exo-) (NEB, M0212) and either Illumina TruSeq DNA LT Kit indexed adapters (Illumina, 20015964) or NEBNext Multiplex Oligos (NEB, E7780S) were employed. Libraries were amplified in PCR reactions for 5–7 cycles using the TruSeq DNA LT kit (Illumina, 15041757) and subjected to SPRI bead size selection before sequencing on either an Illumina HiSeq 4000 or a NovaSeq 6000 using the Paired End 50 bp or 100 bp modules. Two biological replicates were performed for each condition, with the exception of early mitotic exit populations in telophase or cytokinesis.

**ATACseq**

Chromatin accessibility was investigated using omni ATACseq (Corces et al. 2017; Buenrostro et al. 2015) with some modifications. Prometaphase-arrested cells were collected by mitotic shake-off and adherent cells were dissociated using accutase, 5 hours after prometaphase release. 50,000 viable cells were permeabilized in 50ul cold Resuspension Buffer (10mM Tris-HCl pH 7.4, 10mM NaCl, 3mM MgCl2) containing 0.1% NP-40 (MP Biomedicals, 0219859680), 0.1% Tween-20, and 0.01% Digitonin (Promega, G9441) for 3 minutes on ice. Cells were isolated by spinning at 500 RCF for 10 mins at 4°C and the buffer was exchanged by washing once with the detergent-free Resuspension Buffer. Tagmentation was performed in permeabilized cells resuspended in 50ul Tagmentation buffer (Diagenode, C01019043) containing 100nM adaptor-loaded Tn5 transposase (Diagenode, C01070012-30), 0.001% Digitonin, and 0.1% Tween-20 at 37°C for 30 minutes with intermittent mixing at 1000 rpm. The reaction was stopped by addition of Binding Buffer (Qiagen MinElute PCR Kit, 28004) and DNA was purified using the Qiagen MinElute PCR Kit according to the manufacturer's protocol. Purified DNA was eluted in 21 μl of Elution buffer and stored at – 20°C for library preparation. To prepare ATAC-seq libraries for sequencing, custom barcoded primers based on a previous design (Buenrostro et al. 2013) were used in an initial 5 cycle pre-amplification using NEBNext High Fidelity PCR Master Mix (NEB, M0541). The number of PCR cycles for PCR amplification was determined using qPCR. Following PCR-amplification, libraries were purified using SPRI beads, with a sample to bead ratio of 1:1.5. ATAC-seq libraries sequenced on an Illumina NextSeq 2000 machine using the 50 bp paired-end reagents. Two biological replicates were performed for each condition.

**Stable isotope labeling by amino acids in cell culture (SILAC)**

SILAC was performed as described previously (Ong et al. 2002; Ong and Mann 2006). Prior to cell cycle synchronization, DLD-1 RanGAP1-AID and Nup93-AID cells were labeled by culturing over for 5 days in heavy (L-Arginine13C6, 15N4 hydrochloride (Sigma 608033) and (L-Lysine:2HCL 13C6 15N2 (Cambridge Isotope CNLM-291-H)) or light (L-arginine 13C6 hydrochloride (Sigma, 643440) and L-lysine (Sigma L5501)) DMEM for SILAC medium (Thermo Scientific, A33822) with dialyzed FBS (Sigma, F0392). SILAC was maintained throughout the cell synchronization experiment and nuclei were isolated for LC-MS. For each cell line one replicate experiment was performed using “light” control and “heavy” IAA-treated samples and labeling was then reversed for the second biological replicate.

**Nuclear isolation and liquid chromatography mass spectrometry (LC-MS)**

Nuclei were isolated by manual disruption based on a previous protocol (Herrmann et al. 2017) for mass spectrometry analysis. Early G1 cells labeled were collected 5 hours after prometaphase release and washed with PBS. 10 million cells were carefully resuspended with a broad pipette tip in 1 ml of hypotonic buffer (10mM Hepes pH 7.9, 1.5mM MgCl2, 10mM KCl) containing 0.1mM PMSF, 0.5mM dithiothreitol (DTT) (Fisher Scientific, BP172-25), and 1x protease inhibitor cocktail (Thermo Fisher, 78440) and incubated on ice for 25 minutes. Cells were then ruptured by douncing 40x in a pre-chilled homogenizer (tight pestle (B)) and nuclei were isolated by spinning at 1500 x g for 5 minutes at 4°C and resuspended in a 10mM Tris buffer (pH 7.4, +2mM MgCl2). Samples were adjusted to 1x Laemmli buffer (60mM Tris pH6.8, 10% glycerol, 2% SDS, 100mM DTT) and heated to >85°C for 10 minutes prior to gel electrophoresis. Heavy and light samples were mixed at a 1:1 ratio of total protein based on quantification of stained gels (GelCode Blue Safe Protein Stain (Thermo Scientific, 24594)). Final protein gels were stained with GelCode Blue Safe Protein Stain and bands were excised for LC-MS sample processing.

Trypsin digestion was performed overnight at 37°C. After protein digestion and drying by speed vac samples were reconstituted with 25ul of MS solvent (5% acetonitrile and 0.1% Formic acid) and 3.8ul was injected to the Fusion Lumos Orbitrap MS in OTOT mode with a 90 minutes gradient. Peptides were searched against the SwissProt human database in Maxquant and proteins were subject to a 2 peptide cut-off. Two replicates were performed for each condition, reversing the SILAC-labeling, and at 1% FDR more than 3000 proteins were detected in each sample (Summarized in Supplementary tables 4-5). Fold changes and BH-corrected *p*-values comparing control and depleted cell conditions were determined using Q+ analysis by Scaffold.

**Analysis**

**Published tracks and annotations**

Published and publicly available tracks used in this study are summarized in Supplementary Table 3. Epigenetic profiles for H3K4me3, H3K27Ac, CTCF, and H3K27me3 in DLD1 cells (GSE178593: GSM6245909, GSE214012: GSM6597766, GSM6597768 & GSM6597770) and corresponding bigWigs were used throughout the paper for visualization and as indicated to define cCREs. Variant 1 of the MA0139 CTCF motif annotation from the Jaspar database (Rauluseviciute et al. 2024) was used to define convergent extrusion loops.

Publicly available RNAseq data for the DLD1-RanGAP1 cell line (GSE132363: GSM3860900, GSM3860901, GSM3860902) was used to define active genes for visualization and ATACseq analysis. Raw data was processed using the nf-core/rnaseq pipeline, version 3.15.0: https://github.com/nf-core/rnaseq (Ewels et al. 2020). Active genes were defined by having an FPKM > 0 in all three replicates and an FPKM >1 in the pooled dataset.

**Cis Regulatory Elements**

We used a combination of publicly available epigenetic datasets for DLD-1 together with the ATAC-Seq data generated in this study to define cis regulatory elements (CRE) according to the procedure defined by ENCODE (ENCODE Project Consortium et al. 2020). Briefly, we annotated the list of cell-line independent DNAse hypersensitive sites from ENCODE v2 with Z-score transformed ATAC-Seq, H3K4me3 and H9K27ac signal and used the distance to the nearest TSS to assign DHSs to 7 CRE groups: PLS (22555 promoter-like: open, K4me3 enriched, within 200 bp of the nearest TSS), pELS (28863 combined proximal and near enhancer-like: open, K27ac enriched, within 2kb of the nearest TSS), dELS (43807 distal enhancer-like: open, K27ac enriched, at least 2kb away from the nearest TSS), K4me3 (1417 open K4me3: open, K4me3 enriched, at least 200 bp away from the nearest TSS), CTCF (9911 open, CTCF enriched, not enriched in any other marks) and finally, open (43316 open, not enriched in any marks).

**Aggregation stackup analysis**

Stackups demonstrate behavior of a given signal (e.g. H3K4me3 ChIP-Seq) at a set of genomic loci, typically centered at those loci and presented as a heatmap, where each row represents the signal around the individual genomic locus. We take advantage of the Python API (Abdennur 2024) built around UCSC BBI library (Kent et al. 2010) in order to extract a signal stored in a bigWig or bigBed files given a set of same-sized genomic intervals using the *`stackup`* function. We ensure our intervals have the identical size by centering on the loci of interest and providing fixed-length upstream and downstream flanks. For every stackup we also generate a summary signal profile by averaging the stackup across all rows for every column. We used 100 kb for upstream and downstream flank sizes, and aggregated the signal into 100 bins for every row of the stackup, logarithmic color scales were used throughout the stackups with the exception of EV1 profiles and bigBed-derived signals - coverages of “loop” anchors, MCD anchors, etc.

**Hi-C data pre-processing**

Hi-C libraries were processed using the distiller-nf pipeline (Goloborodko et al. 2022), version 0.3.4: paired-end reads were mapped to hg38 human reference genome using *bwa mem* (Li 2013) in a single-sided fashion (-SP); read alignments were parsed and classified into pairwise interactions, or pairs, by *parse* from the *pairtools* package (Open2C et al. 2024b), version 1.0.2, additional *`--walk-policy all`* option was used in order to rescue multi-way interactions (walks); after the removal of duplicates, uniquely mapped and rescued pairs were further filtered according to their alignment quality (MAPQ > 30), and subsequently aggregated into binned contact matrices in the cooler format (Abdennur and Mirny 2020) at 1, 2, 5, 10, 25, 50, 100, 250, 500 and 1000 kb resolutions; contact matrices were normalized using the iterative correction normalization (Imakaev et al. 2012) with the default parameters e.g.: the first 2 diagonals were excluded from balancing at each resolution in order to avoid short-range ligation artifacts; bins with extreme genomic coverage, as detected by MADmax (maximum allowed median absolute deviation) filter (Schwarzer et al. 2017), were masked and excluded from the analysis. See Supplementary Table 1 and Supplementary Figure 1a for summary mapping statistics, as produced by the pairtools module from MultiQC (Ewels et al. 2016). We used insulation tracks and cis-eigvectors to assess the reproducibility of the replicates, which were pooled together as depicted in the Supplementary Figure 1a.

**Extracting Hi-C features**

*Common setting for feature extraction*

In order to extract Hi-C features relevant for our analyses we used the *cooltools* package (Open2c et al. 2024), version 0.7.0, specifically, we leveraged the *cooltools* Python API (application programming interface) for scripting our analyses in the form of Jupyter Notebooks (Kluyver et al. 2016). We used arms of human autosomal chromosomes (chr1-22) as a genome partitioning for the analyses (parameter *“view_df”*) and default parameters in the API function calls unless specified otherwise. Below we provide a brief description of specific functions that were used to extract each individual feature.

*Scaling analysis*

Frequency of interactions as a function of genomic separation (scaling plots, *P(s)*) was calculated using balanced Hi-C data binned at 1kb using *`expected_cis`* function with smoothing in log-space enabled, data for chromosome arms were “aggregated” to generate an average genome-wide “scaling”. The *`gradient`* function from numpy (Harris et al. 2020) package was used to calculate the rate at which interaction frequency is changing with distance, i.e. scaling plot derivatives in log-log space as demonstrated in the “contacts_vs_distance” notebook from the *“open2c_examples”* repository: https://github.com/open2c/open2c_examples.

*Normalization by “expected”*

Most downstream analyses require Hi-C matrices to be “flattened”, i.e. normalized to the decay of interaction frequency with genomic distance. We use the *`expected_cis`* function to calculate such an “expected”. In case of trans, or inter-chromosomal data, matrices are normalized to average levels of inter-chromosomal interactions, calculated using the *`expected_trans`* function. Results of these functions are passed to the downstream analyses when applicable.

*Insulation*

Diamond insulation score (Crane et al. 2015) was calculated using the “insulation” function at 10kb resolution and 100kb averaging window size (size of the insulation diamond).

*Eigenvectors and compartments*

Eigenvector analysis (Imakaev et al. 2012) was performed separately for each chromosome arm using the *`eigs_cis`* function at 10, 25, 50 and 250kb, where gene density was used to “phase” eigenvectors, i.e. eigenvector tracks were “flipped” if they anticorrelated the gene density track. First eigenvectors, EV1 (ones with the highest eigenvalues, that typically correspond to Hi-C compartments) of select samples were saved as bigWig files to use in stackup analysis and visualization.

*Pairwise-class averaging: saddleplots*

To evaluate how different classes of genomic loci interact with each other in 3D, i.e. the saddle-plot-analysis (Imakaev et al. 2012), we used the *`saddle`* function. First, assignment of the classes to genomic loci (e.g. compartment status, cis regulatory element status, etc.) is done on a bin level and passed as a *`track`* parameter to the function. Second, average level of interactions is calculated for each possible combination of classes from the “flattened” contact map (observed-over-expected). Finally, a class-pairwise average interaction matrix is constructed.

Assigning different classes to genomic bins is done in a specific manner: for published DLD1 IPGs (Scelfo et al. 2024), we used the chromatin state assignments directly at 50kb resolution, after merging B2/3 and B4 heterochromatin classes into “B”; cCREs were hierarchically assigned to 10 and 25 kb genomic bins in the following order: promoter-like (PLS), proximal and near enhancer-like (pELS), distal enhancer-like (dELS), K4me3-open, CTCF and finally, open-sites (e.g. if a given 10kb bin contains pELS element, then the entire bin is assigned pELS status, regardless of other elements present in that bin); “continuous” EV1 tracks were digitized into 38 quantiles after excluding 2.5% of extreme EV1 values from each end of the spectrum.

We estimated the strength of A compartment as an enrichment of AA interactions over AB: AA / ((AB+BA)/2), where AA is an average of OE interactions between EV1 quantiles with 20% strongest A-compartment identity, and AB(=BA) is an average of OE interactions between quantiles with 20% strongest A- and B-identities. Similarly for B-compartment, strength was estimated as: BB / ((AB+BA)/2).

*Average pile-up and quantification analysis*

The *`pileup`* function was used, in order to explore the local interaction pattern of a set of 2D genomic features (defined by a pair of genomic locations, e.g. in our case an all-by-all grid of microcompartment domains, with their various subsets and a set of called “loops”). Briefly, local contact maps (normalized by the “expected”) centered on a given feature and with a fixed flank of 100kb size, a “snippet”, were extracted as a stack and then averaged altogether or in groups of features that meet a certain criteria, e.g. in our case - subgroups of the microcompartment grid by genomic distance, subgroups of the grid overlapping other features, like extrusion dots or domains, etc. cis-chromosomal pileups were done on 10kb contact maps, while trans-chromosomal pileups - on 25kb maps unless specified otherwise.

Local interaction patterns, average pileups, were also used to quantify average “strength” of a given genomic feature. Both for “loops” and the grid of microcompartments, “strength” was defined as a ratio between signal in the center of the pileup and the signal in the periphery, specifically, we used 50 by 50 kb window in the center together with 4 60 by 60 kb corners (as a periphery signal) for the grid of MCDs in cis; 25 by 25 kb center (single pixel in the middle) and 4 50 by 50 kb corners (periphery) for the grid of MCDs in trans; 30 by 30 kb center and 4 70 by 70 kb corners (periphery) for “loop”-strength calculations.

*Dot detection (loop calling) and defining extrusion domains*

In order to detect a reference set of extrusion “loops” for our analyses we pooled control Hi-C data for 5hr and 10hr at 10kb resolution (given their similarity, as demonstrated earlier in Supplementary Figure 1c-d) to achieve higher sequencing depth and then used the *“dots”* function to call significantly enriched interactions. We used *`cluster_filtering=False`* and otherwise the default parameters to apply more stringent singleton filtering afterwards. Detected interactions were further filtered to ensure they are compatible with the convergent CTCF/CTCF interaction.

The resulting list of 18615 interactions/loops was also used to define extrusion domains, intuitively, it is the outermost loop from a subgroup of nested loops that defines a domain. Specifically, “loops” were clustered by their anchor “1” in order to group those situated on the same extrusion line, then most upstream anchor “1” of the cluster was used together with the most downstream anchor “2” of the cluster to define such fully inclusive intervals. Nested and “significantly” overlapping (when overlap between intervals is >70% of either of the intervals) intervals from the resulting list were merged yielding the final list of 3401 “domains”.

**Detecting microcompartment domains (MCD)**

We defined MCDs as the genomic loci/anchors that give rise to the strongly interacting off-diagonal rectangular domains that are clearly visible on the 5hr RanGAP1 depletion contact map, and set out to detect them using screening procedure akin to the detection of “dots”: detect enriched pixels that standout relative to the local background, and after grouping them by proximity, detect those groups that continue interacting with others across distances, i.e. those that continue “checkering”. Such “checkering” anchors are the target feature that we aimed to detect in the first place, i.e. the microcompartment domains.

Next we describe specific steps involved in MCD detection. Variable size and shape of observed MCD-MCD interaction domains dictated the choice of convolution kernels that “describe” local vicinity for a given group of pixels (Supplementary Figure 2b): V (“vertical”) kernel is meant to facilitate detection of horizontally elongated domains, whereas H (“horizontal”) - vertically elongated domains, where a square-shaped group of pixels M (“middle”) corresponds to the part of the domain being tested. Convolutional kernels M, V and H were swept across distance decay-corrected contact map (observed-over-expected) at 10kb resolution up to 30MB for computational efficiency, and enriched pixels were selected with a simple thresholding approach: pixels for which the M is at least 2 times as bright as either of the V or H. Density based clustering approach *`OPTICS`* from sklearn package (Pedregosa et al. 2011), version 1.4.1, (Supplementary Figure 2c) was used to filter out singletons and small groups of enriched pixel, and preserve larger more robust groups of enriched pixels (*`min_samples=5`* and *`max_eps=33kb`* parameters were used). Pixels that remain after the clustering step were used to calculate the “coverage” of enriched pixels (or anchor valency) (Supplementary Figure 2d) and finally we apply a 1D peak detection function *`find_peaks`* from scipy package (Virtanen et al. 2020) to the coverage track in order to detect prominent peaks (Supplementary Figure 2e) that are treated as the final MCD anchors. Each anchor is characterized by its footprint interval and a summit - we are using footprints for most of the downstream analysis, except for the average “pileups” and “stackups”, where we use the summits as an MCD genomic coordinate, instead of e.g. the center of the footprint. The exact details of implementation are available on github (https://github.com/dekkerlab/inherited-folding-programs.git).

In total we detected 2105 MCDs in the 5hr RanGAP1 depletion sample and 853 MCDs in the RanGAP1-depleted sample at Cytokinesis, which are >90% contained within the 5hr one. We used 698 Cyto-MCDs that overlap the 5hr ones for downstream analysis and refer to them as the “early” MCDs, whereas the remaining 1407 MCDs from the 5hr samples are referred to as “late”.

**ATAC-seq analysis**

Batch pre-processing of raw ATAC-seq data from control and auxin-treated RanGAP1-AID cells synchronized in prometaphase or released for 5 hours to early G1 was performed using the nf-core/atacseq pipeline, version 2.1.0: https://github.com/nf-core/atacseq (Ewels et al. 2020). Adaptor-trimmed paired-end reads were mapped to the hg38 reference genome using *bwa mem* and filtered according to the standard pipeline for mapping quality, mitochondrial reads, PCR duplicates, and read length (< 2000 bp). Similarity between replicates was confirmed using DESeq2 (Huber 2017) and filtered alignments were merged and de-duplicated across replicates. See Supplementary Table 2, for summary mapping statistics of ATAC-Seq samples generated in this study. Fragment length distributions were determined from bam files using deepTools (Ramírez et al. 2016). Read ends were derived from the filtered alignments (BAM files) and modified to account for tn5 by shifting +/- reads by +4/-5 bp for use in all downstream applications. ATAC-Seq coverage tracks were generated from shifted read ends using BEDtools (Quinlan and Hall 2010), and scaled to 1 million mapped reads.

We used MACS3 (Zhang et al. 2008) to find ATAC-seq peaks of accessibility in the pooled and single replicate datasets using default parameters with a shift/extend of -75/+150. Peaks called in pooled datasets were only considered True when overlapping a peak in each of the two constituent replicates by at least 50%. To compare these pooled peaks between conditions, the union set was merged using bioframe (Open2C et al. 2024a).

Footprinting analysis across all control G1 peaks was performed using TOBIAS (Bentsen et al. 2020). First, ATACseq signal was corrected for Tn5 insertion bias using the filtered alignments (ATACorrect). The corrected signal was then used to calculate a continuous footprinting score (ScoreBigwig), determining local regions of decreased accessibility. Finally, mean transcription factor footprinting scores as well as pairwise differential “binding” scores (BINDetect) were determined for 841 conserved vertebrate motifs (Fornes et al. 2020), across all control G1 peaks or at the subset of G1 peaks overlapping an MCD. The differential binding score for each TF was normalized based on the distribution of scores at all G1 peaks in both cases.

**Supplementary figures**

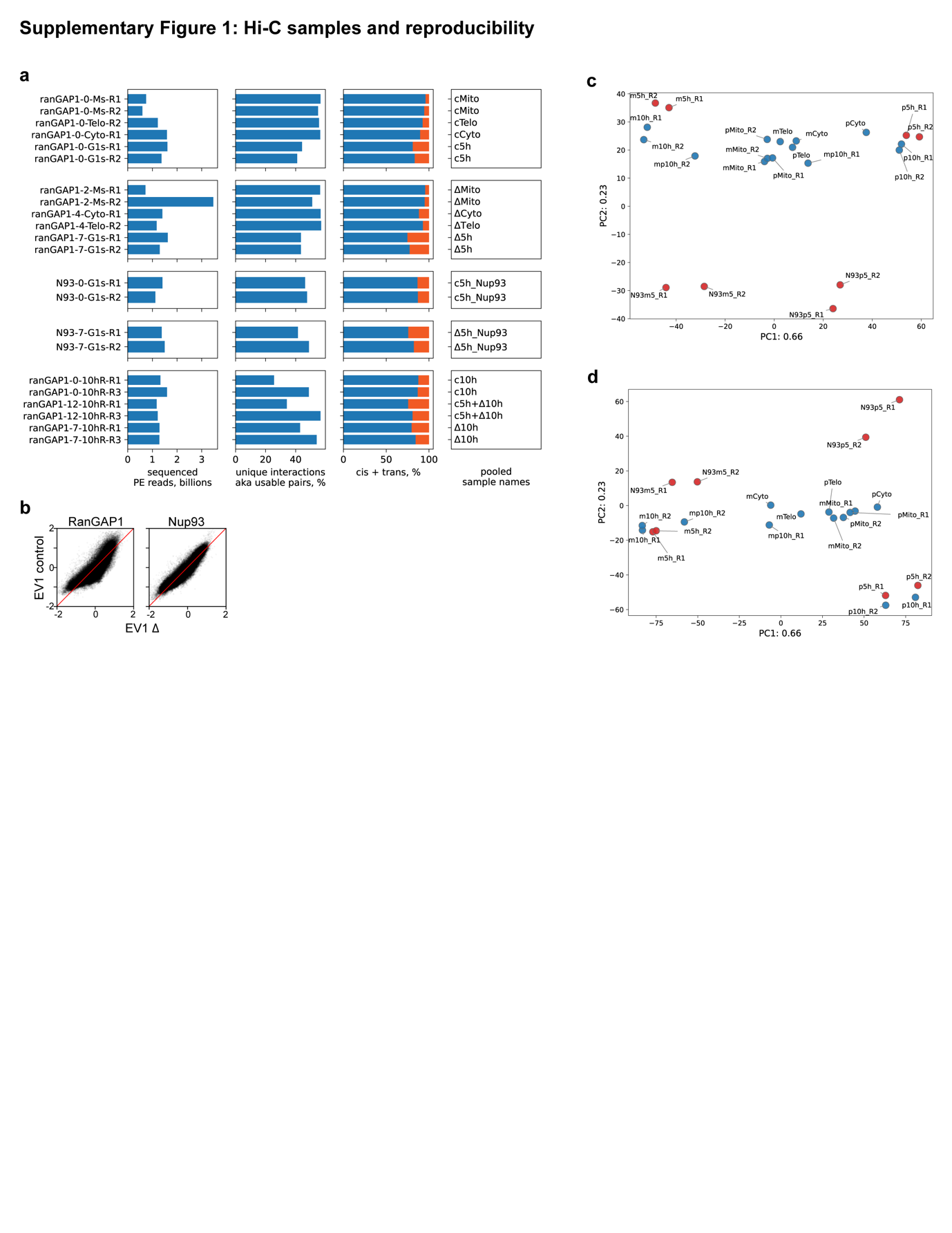

Supplementary Figure1: Hi-C samples and reproducibility

a. Sequencing summary statistics for generated Hi-C samples. Samples are grouped in the following way (vertical arrangement): RanGAP1 control samples across timepoints, RanGAP1 depletion samples across timepoints, Nup93 control sample in G1(5h), Nup93 depletion sample in G1(5h), samples related to the RanGAP1 G1-depletion experiment. The total number of sequenced PE reads is depicted in the first column (in billions), followed by the percentage of uniquely mapped paired end reads (pairs), cis-chromosomal vs. trans-chromosomal percentage breakdown of uniquely mapped pairs, and name of the pooled samples used for all Hi-C related analyses.
b. Comparison of EV1 values from Eigenvector decompositions of 25kb binned Hi-C data from control or Auxin-treated RanGAP1-AID or Nup93-AID cells released to early G1.
c. PCA analysis of the leading eigenvectors (EV1 at 25 kb resolution) across all Hi-C samples. Samples used to define the principal components are depicted in red, the remaining samples were projected onto the derived components. PC1 (explaining >60% of variance) clearly separates depletion samples from the controls (5hr G1), while the projections of earlier time point samples are situated in the middle. Individual replicates of the samples demonstrate concordance according to this metric.
d. PCA analysis of the insulation tracks (at 10 kb resolution, 100 kb diamond size) across all Hi-C samples. Samples used to define the principal components are depicted in red, the remaining samples were projected onto the derived components. PC1 (explaining >60% of variance) clearly separates depletion samples from the controls (5hr G1), while the projections of earlier time point samples are situated in the middle. Individual replicates of the samples demonstrate concordance according to this metric.

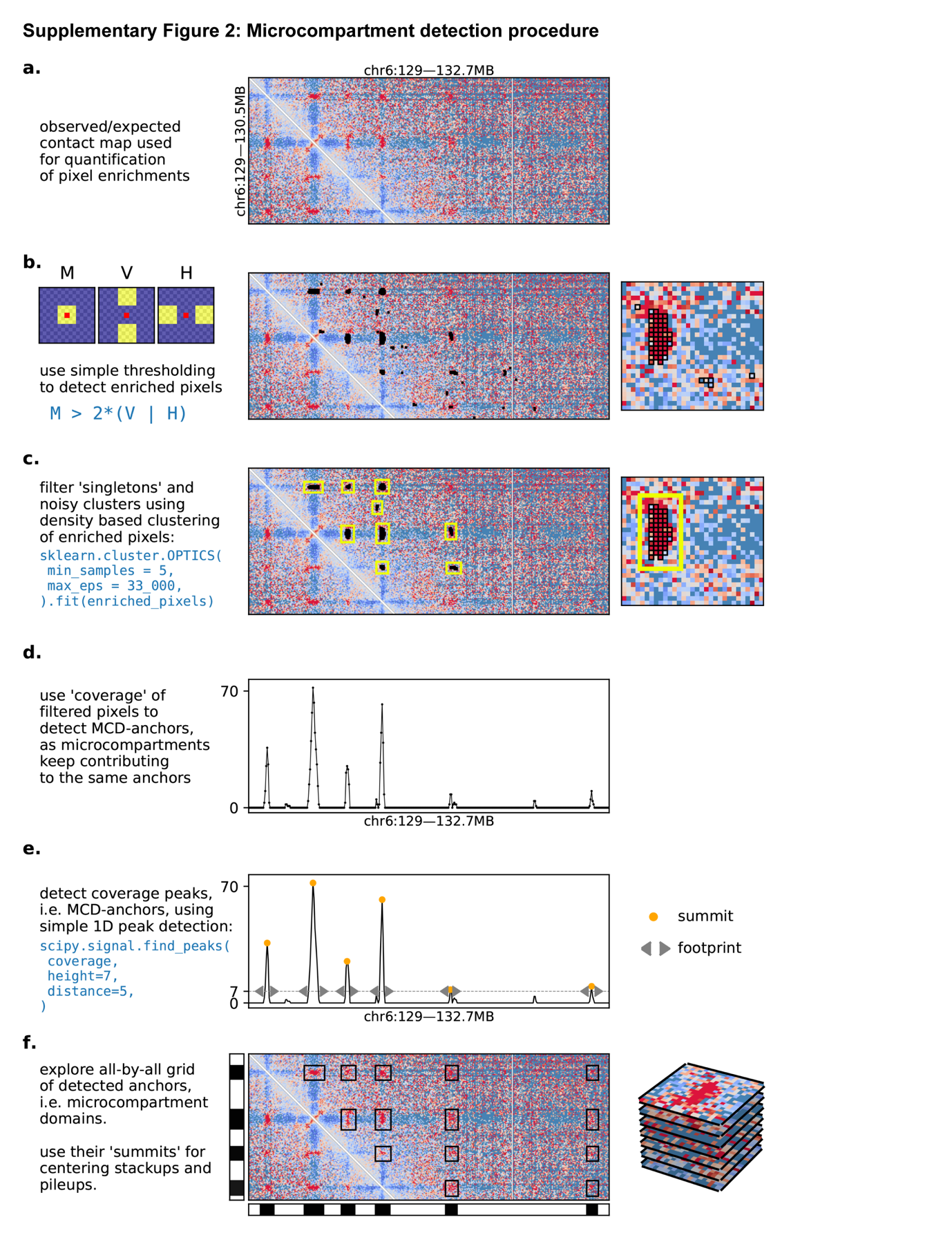

Supplementary Figure 2: Microcompartment detection procedure

a. An example of a “flattened” Hi-C contact map (observed/expected) from the RanGAP1-AID-depleted G1 sample. MCD-MCD interactions are enriched above the expected level and are clearly visible.
b. First step of the MCD detection procedure: Convolution of the contact map with the selected kernels (M, V, H) and thresholding enriched pixels, such that M > 2*(V or H). Enriched pixels are depicted in black, zoom-in is provided on the right.
c. Second step of the MCD detection procedure: Spatial clustering of enriched pixels in order to filter out singleton and other spurious calls. Large clusters that were kept are highlighted with yellow bounding boxes on the selected region of the contact map and the zoom-in (right).
d. Third step of the MCD detection procedure: Enriched pixels from the kept clusters are used to calculate “coverage” (i.e., the number of enriched pixels that overlap a given genomic bin). “Strong” anchors are apparent as prominent peaks on the coverage track for the selected genomic region (center).
e. Fourth step of the MCD detection procedure: 1D peak detection applied to the coverage track results in the detected MCDs shown for the selected genomic region in the center panel. Borders of the peak footprints are depicted in gray, while summits are shown in yellow.
f. Resulting MCDs are used to construct an all-by-all grid or “microcompartment”, as shown in the center panel. Each instance of MCD-MCD interaction is depicted using a black bounding box. A typical way to explore a local pattern of interaction around MCDs is to pile them as shown on the right and to calculate the average signal.
